# Sexual differences in neuronal and synaptic properties across subregions of the mouse insular cortex

**DOI:** 10.1101/2023.10.18.562844

**Authors:** Daniela Iezzi, Alba Cáceres-Rodríguez, Benjamin Strauss, Pascale Chavis, Olivier J. Manzoni

## Abstract

**Background:** The insular cortex (IC) plays a pivotal role in processing interoceptive and emotional information, offering insights into sex differences in behavior and cognition. The IC comprises two distinct subregions: the anterior insular cortex (aIC), that processes emotional and social signals, and the posterior insular cortex (pIC), specialized in interoception and perception of pain. Pyramidal projection neurons within the IC integrate multimodal sensory inputs, influencing behavior and cognition. Despite previous research focusing on neuronal connectivity and transcriptomics, there has been a gap in understanding pyramidal neurons characteristics across subregions and between sexes.

**Methods:** Adult male and female C57Bl/6J mice were sacrificed and tissue containing the IC was collected for ex vivo slice electrophysiology recordings that examined baseline sex differences in synaptic plasticity and transmission within aIC and pIC subregions.

**Results:** Clear differences emerged between aIC and pIC neurons in both males and females: aIC neurons exhibited distinctive features such as larger size, increased hyperpolarization, and a higher rheobase compared to their pIC counterparts. Furthermore, we observed variations in neuronal excitability linked to sex, with male pIC neurons displaying a greater level of excitability than their female counterparts. We also identified region-specific differences in excitatory and inhibitory synaptic activity and the balance between excitation and inhibition in both male and female mice. Adult females demonstrated greater synaptic strength and maximum response in the aIC compared to the pIC. Lastly, synaptic long-term potentiation occurred in both subregions in males but was specific to the aIC in females.

**Conclusions:** We conclude that there are sex differences in synaptic plasticity and excitatory transmission in IC subregions, and that distinct properties of IC pyramidal neurons between sexes could contribute to differences in behavior and cognition between males and females.

**Highlights:**

- Distinctions specific to sex are present within subregions of the insular cortex (IC) in C57Bl/6J mice.
- Pyramidal neurons in the anterior IC (aIC) exhibited larger size and distinct electrical properties. Adult females exhibited stronger synaptic responses in the aIC.
- Conversely, male posterior insular cortex neurons displayed increased excitability.
- Synaptic long-term potentiation was observed in both subregions in males, but it was exclusive to the aIC in females.
- Sex-based variations in various aspects of excitatory transmission within IC subregions could contribute to differences in behavior and cognition between males and females.

**Plain language summary:** This study investigates differences in the insular cortex (IC), a region of the brain responsible for emotions and sensory perceptions, between male and female mice. The IC has two parts: the front (aIC) deals with emotions and social cues, while the back (pIC) is focused on sensing pain and bodily sensations. We examined specific brain cells called pyramidal neurons in both aIC and pIC and discovered noteworthy distinctions between these neurons in adult male and female mice. Firstly, aIC neurons were larger and had unique electrical properties in both male and female mice. Males had more excitable pIC neurons compared to females, indicating that their neurons were more likely to transmit signals. We also explored how these neurons communicate with each other through connections known as synapses. In adult females, the aIC had stronger connections than the pIC. Finally, we observed that specific types of basic synaptic learning occurred exclusively in males in the aIC.

These findings underscore significant disparities in the IC between males and females, offering valuable insights into the potential reasons behind variations in behaviors and emotions between sexes.

## Introduction

The human insula, a deep brain region with extensive connections to various cortical and subcortical regions, is known for its role in processing interoceptive and emotional information ^1^. The insular cortex (IC), is subdivided into two distinct regions: the anterior (aIC) and the posterior IC (pIC), each with unique functions. In humans, the aIC is associated primarily with self-awareness and social emotions like empathy and compassion ^2^, while the pIC handles interoceptive information processing, such as temperature and heart regulation, taste perception, pain processing, and attention to bodily sensations ^3,4^. Animal studies, principally rodents, also corroborate the central role played by the IC in integrating sensory, emotional, and cognitive information. The IC may contribute to sex differences in brain disorders through its involvement in various functions and its differential activation and connectivity patterns between males and females. For example, sex-related IC differences may lead to differences in how emotions are processed and how sensitive individuals are to social emotions, which are mainly associated with the aIC or on pain processing and interoceptive awareness, functions primarily associated with the pIC. These variations could result in differences in how pain is perceived, how stress is responded to, or the degree of attention given to bodily sensations when comparing individuals of different sexes. A current hypothesis is that these differences play a role in the varying prevalence and manifestation of certain disorders between the sexes. Thus, the IC has been implicated in mood disorders, panic disorder, PTSD, obsessive-compulsive disorder, eating disorders, and schizophrenia ^5^ and some of these disorders, such as anxiety and trauma-related disorders, are more common in women than in men ^6–8^. Exploring these sex-related variations in the IC’s subregions is imperative to gain insights into the neural underpinnings of behaviors and cognitive processes specific to each sex. Pyramidal neurons, acting as the primary excitatory neurons within the IC, have a pivotal role in processing interoceptive, emotional, and cognitive information. Their critical function involves transmitting and integrating information within the IC, particularly from the pIC to the aIC, and onwards to output regions, thereby directly influencing both behavior and cognition. Although prior research has extensively examined the connectivity and transcriptomics of these neurons ^9,10^ a comprehensive investigation into the intrinsic passive, active, and synaptic properties of these neurons has been notably lacking. Yet, it is these intrinsic properties that form the very foundation of the IC’s functionalities. To gain a comprehensive understanding of the IC’s operations, it is imperative to systematically compare these intrinsic properties not only between different subregions but also between different sexes. This study focused on examining disparities in cellular and synaptic properties among neurons located in the aIC and pIC regions of mice, taking into consideration variations linked to sex. The results highlighted notable intrinsic dissimilarities between neurons in both the aIC and pIC across both male and female subjects. Moreover, we identified sex-related distinctions in terms of neuronal excitability and synaptic activity. These findings provide valuable insights into the neural underpinnings of sex-related differences in behavior and cognition associated with the IC.

## Materials and methods

### Animals

Animals were treated in compliance with the European Communities Council Directive (86/609/EEC) and the United States NIH Guide for the Care and Use of Laboratory Animals. The French Ethical committee authorized the project APAFIS#18476-2019022510121076 v3. Adult male and female C57BL6/J (12–17 weeks age) were purchased from Charles River and housed in standard wire-topped Plexiglas cages (42□×□27□x□14□cm), in a temperature and humidity-controlled condition (i.e., temperature 21□±□1□°C, 60□±□10% relative humidity and 12□h light/dark cycles). Food and water were available ad *libitum*.

### Slice preparation

Adult male and female mice (PND 90-120) were deeply anesthetized with isoflurane and sacrificed according to institutional regulations. The brain was sliced (300□μm) in the coronal plane with a vibratome (Integraslice, Campden Instruments) in a sucrose-based solution at 4°C (87 mM NaCl, 75 mM sucrose, 25 mM glucose, 2.5 mM KCl, 4 mM MgCl2, 0.5 mM CaCl2, 23 mM NaHCO3, and 1.25 mM NaH2PO4). Immediately after cutting, slices containing anterior or posterior IC were stored for 30 h at 32°C in a low-calcium artificial CSF (ACSF) that contained the following: 130 mM NaCl, 11 mM glucose, 2.5 mM KCl, 2.4 mM MgCl2, 1.2 mM CaCl2, 23 mM NaHCO3, and 1.2 mM NaH2PO4, and were equilibrated with 95% O2/5% CO2 and then at room temperature until the time of recording. During the recording, slices were placed in the recording chamber and continuously perfused at 2 ml/min with warm (32°-34°C) low Ca2+ solution.

### Electrophysiology

Whole-cell patch clamp recordings were made from the soma of layer V pyramidal anterior or posterior IC neurons. The latter were visualized under a differential interference contrast microscope using an upright microscope with infrared illumination (Olympus, France). For current-clamp experiments and voltage clamp recording, patch pipettes were filled with an intracellular solution containing (in mM): K^+^ gluconate (145 K^+^ gluconate, 3 NaCl, 1 MgCl_2_, 1 EGTA, 0.3 CaCl_2_, 2 Na^2+^ ATP, 0.3 Na^+^ GTP, and 0.2 cAMP, buffered with 10 HEPES). The pH was adjusted to 7.25 and osmolarity to 290 – 300 mOsm. Electrode resistance was 2 – 4 MΩ. Access resistance compensation was not used, and acceptable access resistance was <30 MΩ. The potential reference of the amplifier was adjusted to zero before breaking into the cell. Cells were held at –70 mV. Current-voltage (I-V) curves were made by a series of hyperpolarizing to depolarizing current steps immediately after breaking into the cell. To determine rheobase a series of depolarizing current steps was applied. Spontaneous EPSCs (sEPSCs) were recorded at −70 mV and isolated by using the GABA_A_ receptor blocker gabazine 10 mM (SR 95531 hydrobromide; Tocris).

When inhibitory postsynaptic currents (IPSCs) were recorded, the recording pipettes were filled with a high-chloride solution of the following composition (in mM): 140 KCl, 1.6 MgCl_2_, 2.5 MgATP, 0.5 NaGTP, 2 EGTA, 10 HEPES. The pH solution was adjusted to 7.25-7.3 and osmolarity to 280 – 300 mOsm. Electrode resistance was 3–4 MΩ. Spontaneous IPSCs (sIPSCs) were recorded at −70mV in presence of 20□μM CNQX (6-Cyano-7-nitroquinoxaline-2,3-dione disodium, an AMPA receptor antagonist, Tocris) and L-APV 50 μM (DL-2-Amino-5-phosphonopentanoic acid, a selective NMDA receptor antagonist, Tocris).

For extracellular field potential recordings, both stimulating and recording electrode were positioned in the layer V of a/pIC. Both fEPSP area and amplitude were analyzed. Stimulation was performed with a glass electrode filled with aCSF and the stimulus intensity was adjusted ∼60% of maximal intensity after performing an input–output curve in presence of gabazine (10 mM). For LTP experiments, baseline stimulation frequency was set at 0.1 Hz and plasticity was induced by a high-frequency stimulation (HFS; stimuli were applied at 100□Hz for 1□s, repeated 3 times at 10□s interval). The glutamatergic nature of the field EPSP (fEPSP) was systematically confirmed at the end of the experiments using the ionotropic glutamate receptor antagonist CNQX (20 mM), which specifically blocked the synaptic component without altering the non-synaptic.

Data was recorded in current clamp with an Axopatch-200B amplifier, low pass filtered at 2 kHz, digitized (10 kHz, DigiData 1440A, Axon Instruments), collected and analyzed using Clampex 10.7 (Molecular Device).

### Data analysis and statistics

except for Principal Component Analysis (PCA, see below), data were analysed off-line with Clampfit 10.7 (Molecular Devices, Sunnyvale, CA, USA) and AxoGraph X. Graphs and Figure layouts were generated with GraphPad Prism 7.0. Datasets were tested for the normality (D’Agostino-Pearson and Shapiro-Wilk) and outliers (ROUT test) before running parametric tests. Statistical significance of difference between means was assessed with two-way ANOVA followed by Sidak’s multiple comparison post hoc tests as indicated in figure legends. When achieved, the significance was expressed as exactly *p*-value in the figures. The experimental results are described qualitatively in the main text, whereas experimental and statistical details, including sample size (n/N = cells/animals), statistical test, *p*-value, main effects and interactions, are reported in Figure legends or in Supplementary Tables. Quantitative data are presented as Box and whisker plots, reporting median, min and max values, and superimposed scatter plots to show individual data points.

Membrane capacitance (Cm) was estimated by integrating the capacitive current evoked by a −2 mV pulse, whereas the membrane resistance was estimated from the I-V curve around resting membrane potential. The latter (RMP) was measured immediately after whole-cell formation during the current-clamp protocol. The input-output curve was made by measuring the number of action potentials elicited by depolarizing current steps of increasing amplitude, while to determine rheobase a series of depolarizing 10 pA current steps was applied. The frequency and amplitude of sE/IPSCs were analysed with Axograph X using a double exponential template: f(t) = exp(-t/rise) + exp(-t/decay) (rise = 0.5 ms and decay = 3 ms; rise = 0.2 ms and decay = 10 ms, respectively). The detection threshold for the events was set at 3 times the baseline noise SD, whereas the one for the amplitude detection was set at -7 pA. Total charge was calculated by summing the charge transfer of all individual events (sEPSCs or sIPSCs) detected over a 6 min acquisition period for each neuron.

For field recording experiments, the magnitude of plasticity was calculated at 0–10 and 20– 30□min after induction as percentage of baseline responses.

Considering the parameters reported in the evaluation of intrinsic and synaptic transmission properties, PCA was computed using “The FactoMineR package”. Intrinsic properties of layer 5 pyramidal neurons were analysed via PCA with membrane capacitance, rheobase, resting membrane potentials, neuronal excitabilities and voltage membrane response to a different injected current steps as quantitative variables and individual cells as individuals. Missing values were imputed by the mean of the variable. Supplementary qualitive variables were the two insular cortexes (anterior and posterior, two modalities), sex (2 modalities) and group (4 modalities). The cumulative relative contribution of PCs against the variance, the contribution and correlation were investigated.

## Results

### Intrinsic properties of pyramidal neurons in the anterior and posterior IC of both sexes

In the rostro-caudal axis, the IC is divided into two regions: the anterior insula (aIC) and the posterior insula (pIC). While the distinct reciprocal connections of these regions with cortical and subcortical brain areas have been well-characterized, less is known about their intrinsic properties. Thus, electrophysiological characterizations of pyramidal neurons in both aIC and pIC were conducted in male and female subjects: layer V pyramidal neurons were whole-cell patch clamped in acute coronal slices from mouse aged between 90 to 120 postnatal days (PNDs) (**Fig. 1A-B**).

**Figure 1:**
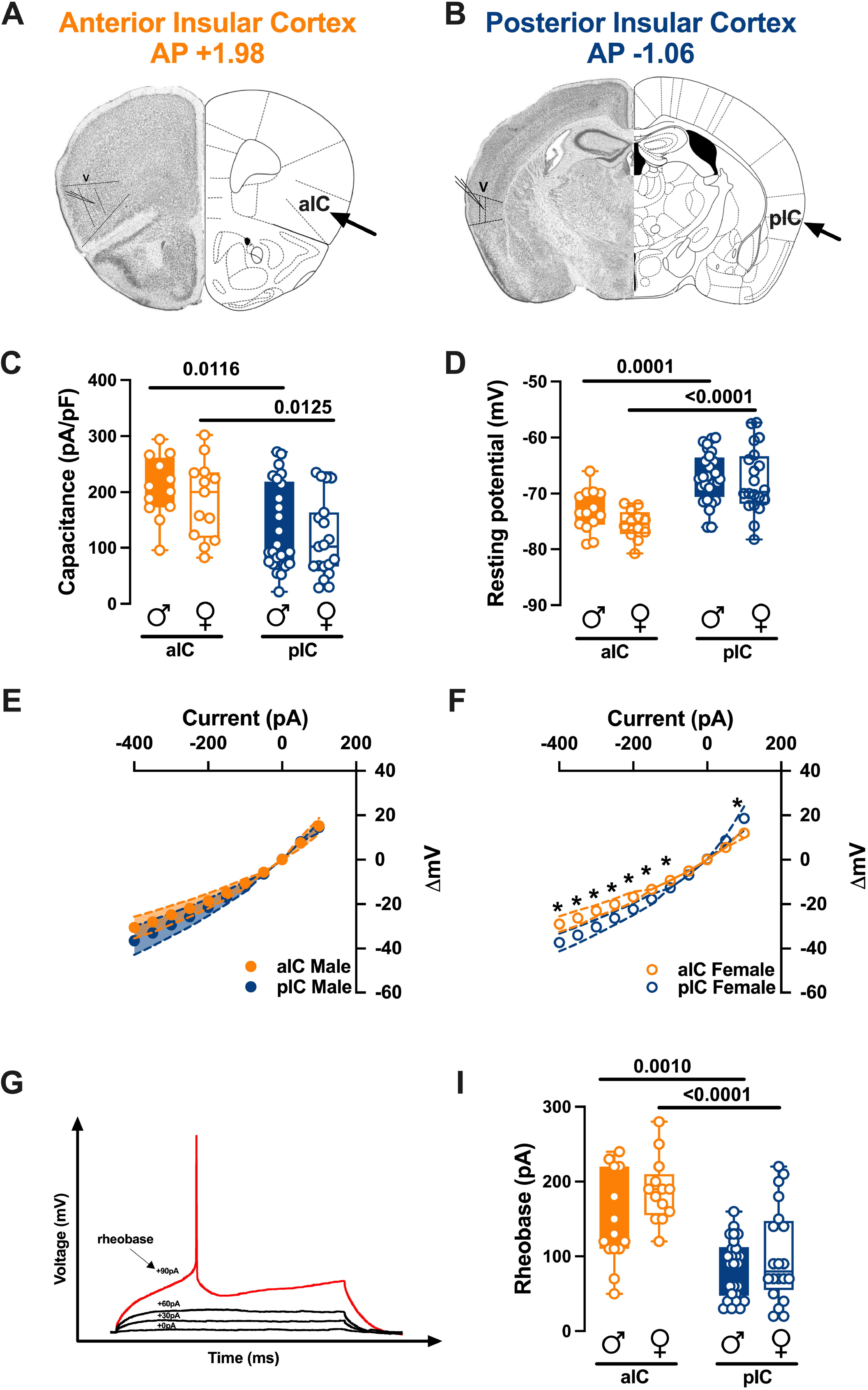
Subregional intrinsic properties of anterior and posterior insular cortex pyramidal neurons in males and females. **(A-B)** Representative coronal sections of the brain from the anterior (Bregma +1.98) and posterior (Bregma -1.06) regions of the insular cortex (IC) showing recording sites of layer V pyramidal neurons. **(C-D)** Quantitative analysis of passive and active membrane properties revealed that anterior IC (aIC) pyramidal neurons are significantly larger (i.e., larger capacitance) and hyperpolarized compared to posterior IC (pIC) neurons, and that this difference is present in both male and female mouse. **(E-F)** Subregional differences in the current-voltage relationship between aIC and pIC pyramidal neurons were found in response to current injection steps of 50 pA, ranging from -400 pA to +50 pA, in adult females **(E)**, but not in males **(F)**. **(G)** An example of an action potential (AP) evoked by increasing current steps is illustrated, with the rheobase indicated. **(I)** The results indicated that the minimal current required to trigger an AP (rheobase) was significantly higher in aIC compared to pIC pyramidal neurons. Data are presented as box-and-whisker plots (minimum, maximum, median) for **(C-D-I)**, and as mean ± SEM in XY plots for **(E-F)**. Two-way ANOVA followed by Šídák’s multiple comparison test was performed for **(C-D-I)**, and Mann-Whitney U test was applied for **(E-F)**. P-values <0.05 are depicted in the graphs. The sample sizes for aIC male and female were 14/10 and 13/6, respectively, and for pIC male and female were 28/15 and 20/12, respectively.

There were no significant differences in passive or active membrane properties when comparing males and females within each IC region. However, substantial differences were observed between the anterior and posterior IC. Specifically, pyramidal neurons in the aIC were notably bigger and hyperpolarized compared to those in the pIC (**Fig. 1C-D**, **Table 1**). Furthermore, aIC exhibited a higher rheobase compared to pIC neurons in both males and females, consistent with their hyperpolarized resting potential (**Fig. 1I**, **Table 1**).

**Table 1.** Active and passive membrane properties of aIC and pIC pyramidal neurons in both sexes.

| Misure | Area | Sex | Median | MAX | MIN | n/N | Two-way Anova<br>Main effects and<br>interactions | Multiple<br>comparison<br>(Šidák's post<br>hoc test) |
| --- | --- | --- | --- | --- | --- | --- | --- | --- |
| **Cm (pA/pF)** | aIC | M | 205.9 | 294 | 95.67 | 14/10 | Interaction:<br>p = 0.9586<br>Area:<br>F (1.65) = 15.93<br>p = 0.0002<br>Sex:<br>F (1.65) = 1.323<br>p = 0.2543 | aIC Male vs<br>pIC Male<br>p = 0.0103 |
|  |  | F | 200.2 | 302.1 | 82.38 | 13/6 |  | aIC Female vs<br>pIC Female<br>p = 0.0153 |
|  | pIC | M | 92.73 | 271.8 | 21.66 | 28/15 |  | aIC Male vs<br>aIC Female<br>p = 0.7007 |
|  |  | F | 102.1 | 235.2 | 28.54 | 20/12 |  | pIC Male vs<br>pIC Female<br>p = 0.6001 |
| **Vr (mV)** | aIC | M | -73.45 | -66 | -79.07 | 14/10 | Interaction:<br>p = 0.4244<br>Area:<br>F (1.70) = 39.08.<br>p = <0.000<br>Sex:<br>F (1.70) = 1.791.<br>p = 0.1852 | aIC Male vs<br>pIC Male<br>p = 0.0002 |
|  |  | F | -75.34 | -71.75 | -80.78 | 13/6 |  | aIC Female vs<br>pIC Female<br>p = <0.0001 |
|  | pIC | M | -67.19 | -60 | -76 | 28/15 |  | aIC Male vs<br>aIC Female<br>p = 0.3378 |
|  |  | F | -69.69 | -57.34 | -78.22 | 20/12 |  | pIC Male vs<br>pIC Female<br>p = 0.8815 |
| **Rh (pA)** | aIC | M | 125 | 240 | 50 | 14/10 | Interaction:<br>p = 0.3339<br>Area:<br>F (1.69) = 35.52 p<br>= <0.0001<br>Sex:<br>F (1.69) = 5.301<br>p = 0.0243 | aIC Male vs<br>pIC Male<br>p = 0.0010 |
|  |  | F | 190 | 280 | 120 | 13/6 |  | aIC Female vs<br>pIC Female<br>p = <0.0001 |
|  | pIC | M | 90 | 160 | 30 | 28/15 |  | aIC Male vs<br>aIC Female<br>p = 0.0828 |
|  |  | F | 80 | 220 | 20 | 20/12 |  | pIC Male vs<br>pIC Female<br>p = 0.4825 |

When comparing the membrane voltage response to hyperpolarizing current steps, we observed that in males, there were no significant differences between aIC and pIC (**Fig. 1E, Table 1A**). However, in females, the voltage response of aIC was significantly lower compared to pIC (**Fig. 1F, Table 1A**). Finally, no sex differences were observed when comparing the voltage membrane responses in both cortexes (**Supplementary Fig. 1A-B, Suppl. Table 1**). Principal Component Analysis (PCA, **Fig.3**, **Table 3**) with membrane capacitance, rheobase, resting membrane potentials, neuronal excitabilities, and voltage membrane response to a different injected current steps as quantitative variables, indicated that subregion if the principal contributor to the dataset variance when all groups are considered (**Fig. 3B**).

**Table 1A.** Voltage membrane response per current steps in aIC and pIC pyramidal neurons in both sexes.

| Measure<br>(unit) | Condition | Injected<br>current<br>(pA) | aIC |  |  | pIC |  |  | *p value*<br>*Multiple*<br>*Mann-*<br>*Whitney*<br>*Unpaired t*<br>*test* |
| --- | --- | --- | --- | --- | --- | --- | --- | --- | --- |
|  |  |  | Mean | SEM | n/N | Mean | SEM | n/N |  |
| Voltage<br>changes per<br>current step<br>($\Delta V_m$) | Male | -400 | -30.53 | 2.28 | 14/10 | -36.5 | 3.14 | 28/15 | 0.34 |
|  |  | -350 | -27.98 | 2.18 |  | -32.99 | 2.83 |  | 0.39 |
|  |  | -300 | -24.89 | 1.98 |  | -29.29 | 2.51 |  | 0.35 |
|  |  | -250 | -21.98 | 1.8 |  | -25.65 | 2.22 |  | 0.37 |
|  |  | -200 | -18.51 | 1.6 |  | -21.46 | 1.85 |  | 0.39 |
|  |  | -150 | -14.77 | 1.31 |  | -16.94 | 1.5 |  | 0.47 |
|  |  | -100 | -10.62 | 1 |  | -12.15 | 1.13 |  | 0.59 |
|  |  | -50 | -5.68 | 0.58 |  | -6.49 | 0.63 |  | 0.47 |
|  |  | 0 | 0.05 | 0.04 |  | 0.16 | 0.06 |  | 0.07 |
|  |  | 50 | 7.62 | 1.21 |  | 8.13 | 0.93 |  | 0.66 |
|  |  | 100 | 15.21 | 1.68 |  | 14.58 | 1.19 |  | 0.88 |
|  | Female | -400 | -28.93 | 1.65 | 13/6 | -37.23 | 1.94 | 20/12 | 0.003 |
|  |  | -350 | -26.19 | 1.6 |  | -33.91 | 1.78 |  | 0.001 |
|  |  | -300 | -22.97 | 1.21 |  | -30.23 | 1.61 |  | 0.002 |
|  |  | -250 | -20.08 | 1.2 |  | -26.34 | 1.43 |  | 0.003 |
|  |  | -200 | -16.75 | 1.1 |  | -22.33 | 1.28 |  | 0.003 |
|  |  | -150 | -13.31 | 0.86 |  | -17.74 | 1.1 |  | 0.006 |
|  |  | -100 | -9.3 | 0.62 |  | -12.68 | 0.86 |  | 0.01 |
|  |  | -50 | -5.01 | 0.38 |  | -6.85 | 0.55 |  | 0.02 |
|  |  | 0 | 0.09 | 0.08 |  | 0.21 | 0.15 |  | 0.34 |
|  |  | 50 | 5.52 | 0.47 |  | 8.74 | 0.97 |  | 0.07 |
|  |  | 100 | 11.92 | 0.99 |  | 18.63 | 2.57 |  | 0.02 |

**Table 2.** Intrinsic excitability of aIC and pIC pyramidal neurons in both sexes.

| Measure<br>(unit) | Sex | Injected<br>current<br>(pA) | aIC |  |  | pIC |  |  | *p value*<br>*Multiple*<br>*Mann-*<br>*Whitney*<br>*Unpaired t*<br>*test* |
| --- | --- | --- | --- | --- | --- | --- | --- | --- | --- |
|  |  |  | Mean | SEM | n/N | Mean | SEM | n/N |  |
| Number of<br>APs | Male | 0 | 0 | 0 | 14/10 | 0 | 0 | 20/15 | >0.999999 |
|  |  | 50 | 0.07 | 0.07 |  | 1.05 | 0.42 |  | 0.12 |
|  |  | 100 | 1.36 | 0.73 |  | 4.75 | 1.18 |  | 0.09 |
|  |  | 150 | 3.93 | 0.96 |  | 9.95 | 1.57 |  | 0.01 |
|  |  | 200 | 6.36 | 1.51 |  | 14.95 | 1.78 |  | 0.003 |
|  |  | 250 | 9 | 1.61 |  | 19.15 | 1.95 |  | 0.001 |
|  |  | 300 | 12.07 | 1.56 |  | 21.95 | 1.97 |  | 0.001 |
|  |  | 350 | 14.64 | 1.41 |  | 23.25 | 2.14 |  | 0.008 |
|  |  | 400 | 17 | 1.28 |  | 23.45 | 2.14 |  | 0.02 |
|  |  | 450 | 18.43 | 1.2 |  | 23 | 2.42 |  | 0.07 |
|  |  | 500 | 19.5 | 1.13 |  | 22.85 | 2.45 |  | 0.11 |
|  |  | 550 | 21 | 1.15 |  | 22.85 | 2.71 |  | 0.43 |
|  |  | 600 | 21 | 1.26 |  | 19.35 | 3.14 |  | 0.67 |
|  | Female | 0 | 0 | 0 | 14/6 | 0 | 0 | 16/12 | >0.999999 |
|  |  | 50 | 0 | 0 |  | 0.94 | 0.48 |  | 0.10 |
|  |  | 100 | 0.14 | 0.1 |  | 3.13 | 1.03 |  | 0.02 |
|  |  | 150 | 1.43 | 0.6 |  | 6.5 | 1.44 |  | 0.008 |
|  |  | 200 | 3.79 | 1.06 |  | 9.81 | 1.64 |  | 0.008 |
|  |  | 250 | 6.86 | 1.23 |  | 13.25 | 1.47 |  | 0.005 |
|  |  | 300 | 9.79 | 1.31 |  | 15.88 | 1.33 |  | 0.002 |
|  |  | 350 | 12.29 | 1.41 |  | 18.06 | 1.27 |  | 0.005 |
|  |  | 400 | 14 | 1.51 |  | 19.63 | 1.2 |  | 0.006 |
|  |  | 450 | 15.86 | 1.61 |  | 20.94 | 1.09 |  | 0.02 |
|  |  | 500 | 17.5 | 1.44 |  | 21.38 | 0.9 |  | 0.04 |

**Table 3.**
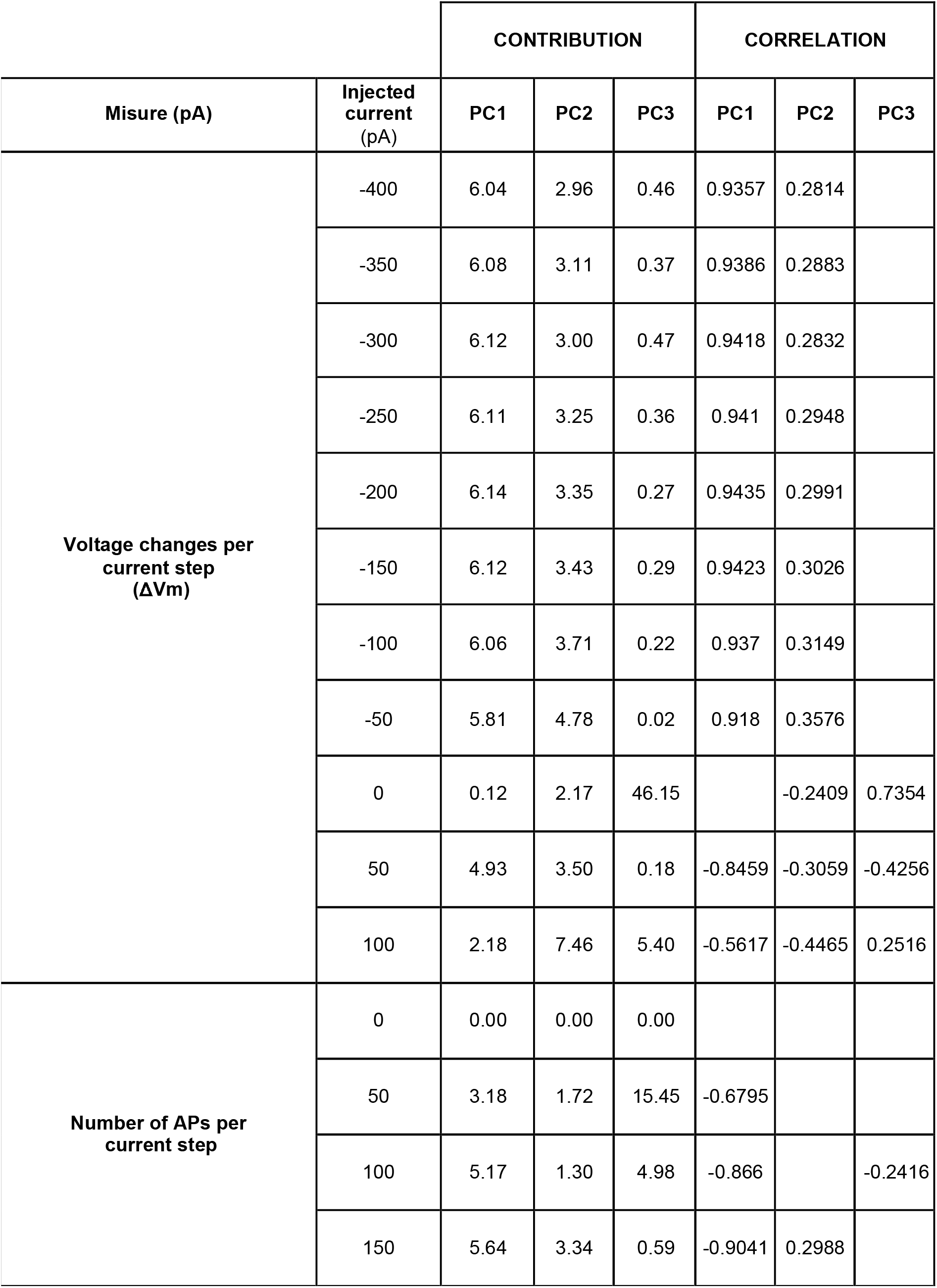
Principal Component Analysis (PCA) of IC pyramidal neurons intrinsic properties.

### Excitability differences in anterior and posterior insular cortex pyramidal neurons by sex and subregion

The observed differences in passive and active membrane properties between the aIC and pIC suggest distinct levels of excitability of the principal projecting neurons in these two subregions.

To assess the intrinsic excitability of IC principal neurons, we recorded their membrane response profiles in reaction to a series of somatic current steps. Both male and female mouse exhibited significant differences in membrane profiles between the aIC and pIC, indicating a higher excitability of pyramidal neurons in the posterior IC (**Fig. 2A-B**, **Table 2**). However, when comparing sexes, pyramidal neurons showed similar excitability (**Supplementary. Fig. 1C-D, Suppl. Table 1**). Furthermore, within the “physiological range”, neurons in the aIC fired less frequently than those in the pIC, independently of sex. Notably, among males, pyramidal neurons in the posterior IC fired more frequently than in females (**Fig. 2D**, **Table 1**). Consistent with these intrinsic passive properties, PCA revealed that the two distinct subregions were the primary contributors to the variance in this dataset (**see Fig. 3B**)

**Figure 2:**
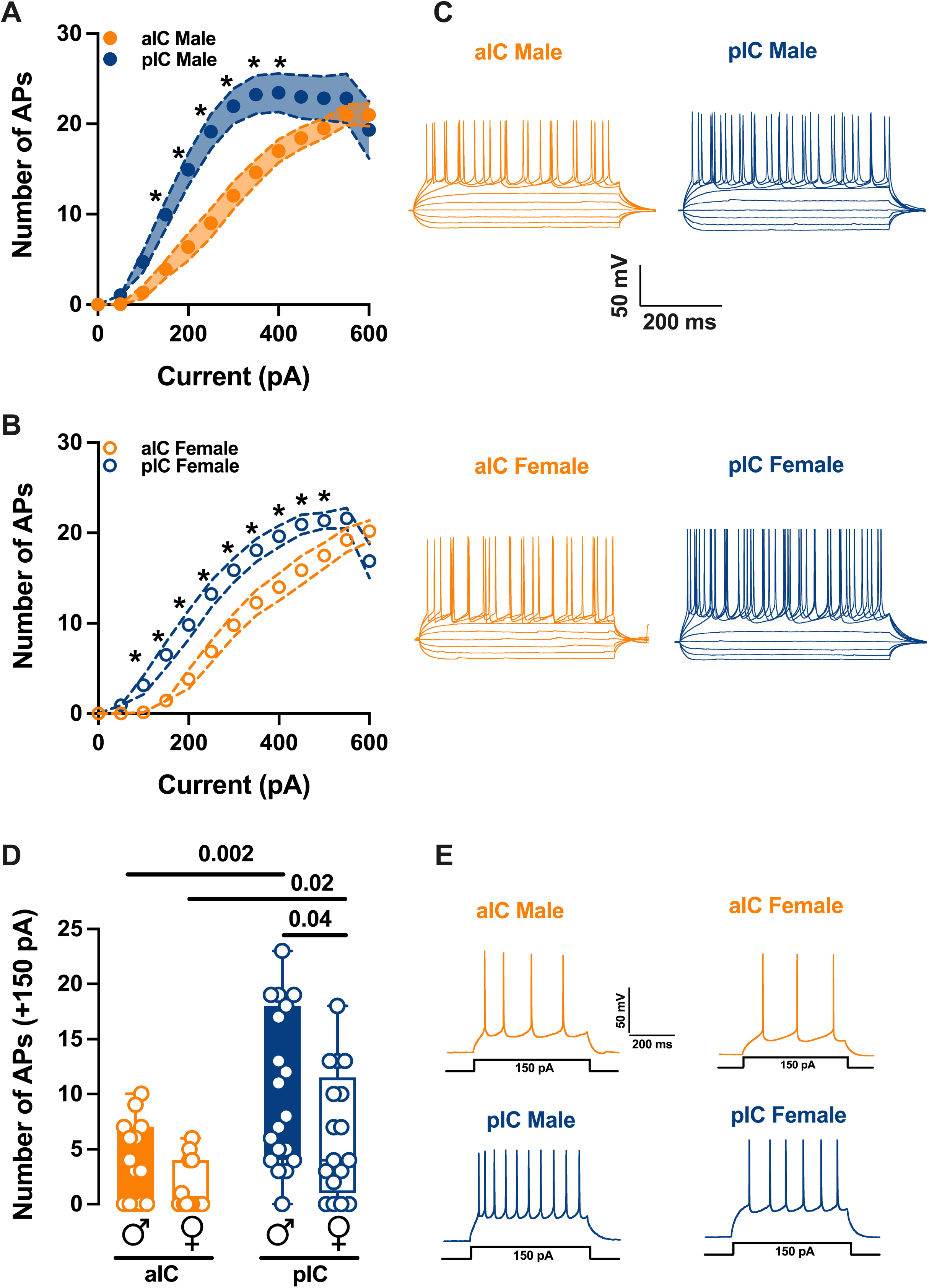
Sex and subregional differences in the excitability of insular cortex pyramidal neurons. **(A-B)** As indicated by the response of action potentials to progressive depolarizing current injection, pyramidal neurons exhibit a notably higher level of excitability in the pIC compared to the aIC, and this trend is consistent across both sexes. (500 ms, ranging from 0 pA to 600 pA in 50 pA steps). **(C)** Representative voltage traces evoked by the injection of hyper- and depolarizing current steps (500 ms, ranging from -150 pA to 250 pA in 50 pA steps) in both anterior and posterior IC pyramidal neurons of both sexes. **(D)** During physiological depolarizing current stages, neurons from the aIC displayed less frequent firing compared to those from the pIC. Additionally, male pIC neurons showed a higher level of excitability than those of their female counterparts. **(E)** An example of firing patterns triggered by the injection of +150 pA depolarizing currents for each group is provided. Data are presented as mean ± SEM in XY plots for **(A-B)**, and as box-and-whisker plots (minimum, maximum, median) for **(C)**. Mann-Whitney U test was applied for **(A-B)**, and two-way ANOVA followed by Šídák’s multiple comparison test was performed for **(C)**. P-values <0.05 are depicted in the graphs. The sample sizes for aIC male and female were 14/10 and 14/6, respectively, and for pIC male and female were 20/15 and 16/12, respectively.

**Figure 3:**
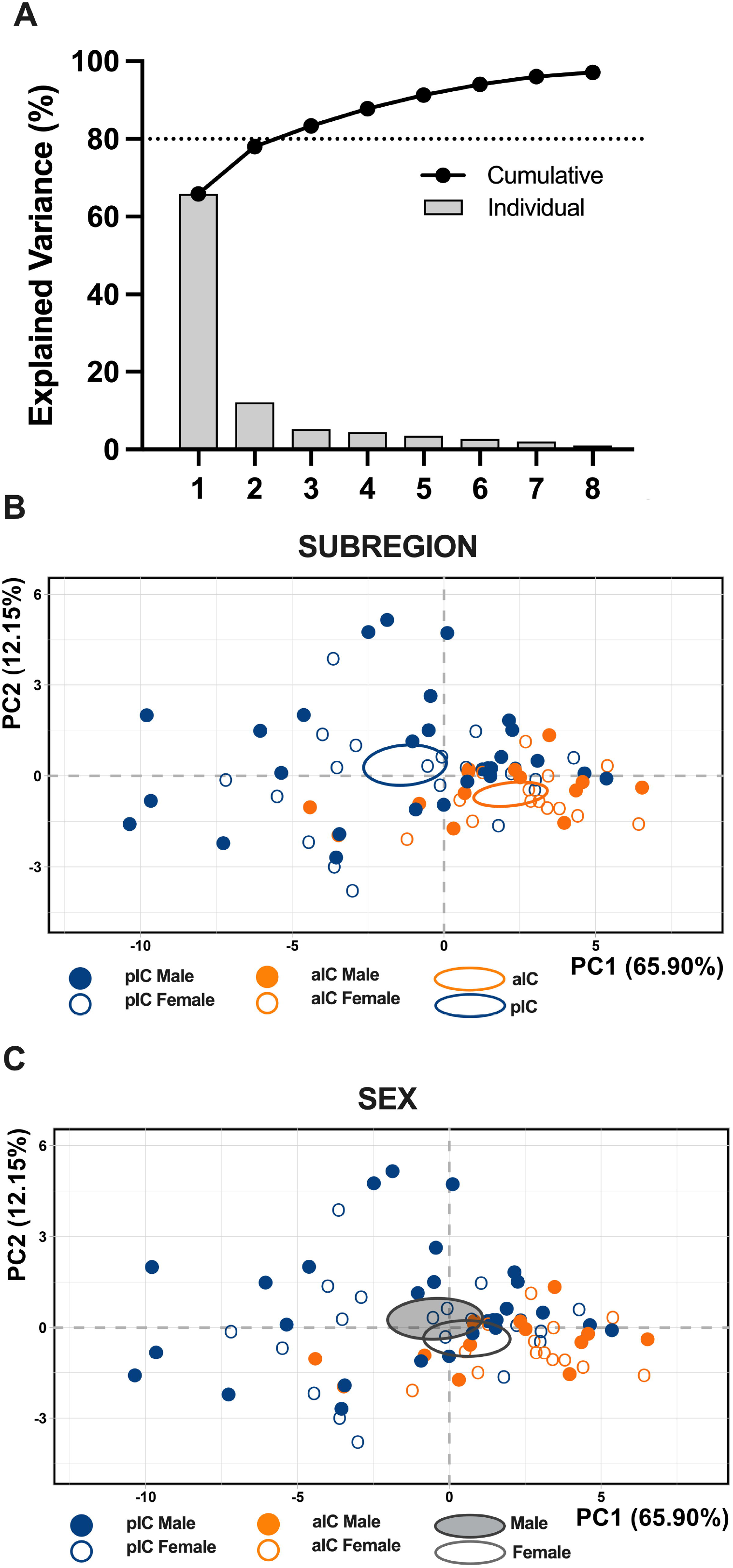
Principal Component Analysis (PCA) of intrinsic properties indicating that subregion is the main factor contributing to the variance in the dataset. This analysis was carried out using membrane capacitance, rheobase, resting membrane potentials, neuronal excitabilities, and the voltage membrane’s response to varying injected current steps as quantitative variables. **(A)** Plotting the percentage of explained variance by each PC (histogram) reveals that most of the data set’s variance is explained by PC1 (65.90%), followed by PC2 (12.15%) and PC3 (5.33%). Black dots represent the cumulative percentage of explained variance. **(B-C)** PCA graphs of individuals were built with PC1 and PC2 which together explained more then 80% of the variance (see A). Small dots represent individuals colored according to their belonging to one the following qualitative supplementary variables: subregion (top), sex (below). Ellipses represent the barycenter of individuals (i.e., mean) for each category. **(B)** PCA show that the subregion is the major contributor in the overall variance. **(C)** In contrast PCA showed that in both insular cortices male and female largely overlap.

### The Excitatory/ Inhibitory balance of across sex and IC subregions

Alterations in excitatory (glutamatergic)/ inhibitory (GABAergic) neurotransmission in the insular cortex have been proposed to play a causal role in the development of chronic pain state and perception of painful stimuli ^11^. However, the E/I balance of IC subregions has not been characterized yet. First, we quantified the total charge transferred from whole-cell recorded spontaneous AMPA-mediated EPSCs (sEPSCs; **Fig. 4A**) and GABA-mediated IPSCs (sIPSCs; **Fig. 4B**) a parameter which accounts for both frequency and amplitude of spontaneous events. The total charge transfer of sEPSCs in aIC was greater than that of the pIC in male only (**Fig 4C**, **Table 4**). The total charge transfer of sIPSCs was similar across sexes and subregions (**Fig. 4D**, **Table 4**). Subsequently, we compared the relative distribution of sEPSCs and sIPSCs total charge transfer ^12,13^. Interestingly, in both sexes and subregions, sIPSCs were predominant (**Fig. 4E-H**) as show by the right-shift in the sIPSCs cumulative distribution, indicating that the E/I balance is shifted towards inhibition (**Fig. 4E-H**, Dot plots). Although the E/I balance in the IC is mainly influenced by inhibition, it is important to note that in both male and female the percentage of inhibition is larger in the pIC than in the aIC (**Fig. 4E-H**, Pie chart).

**Figure 4:**
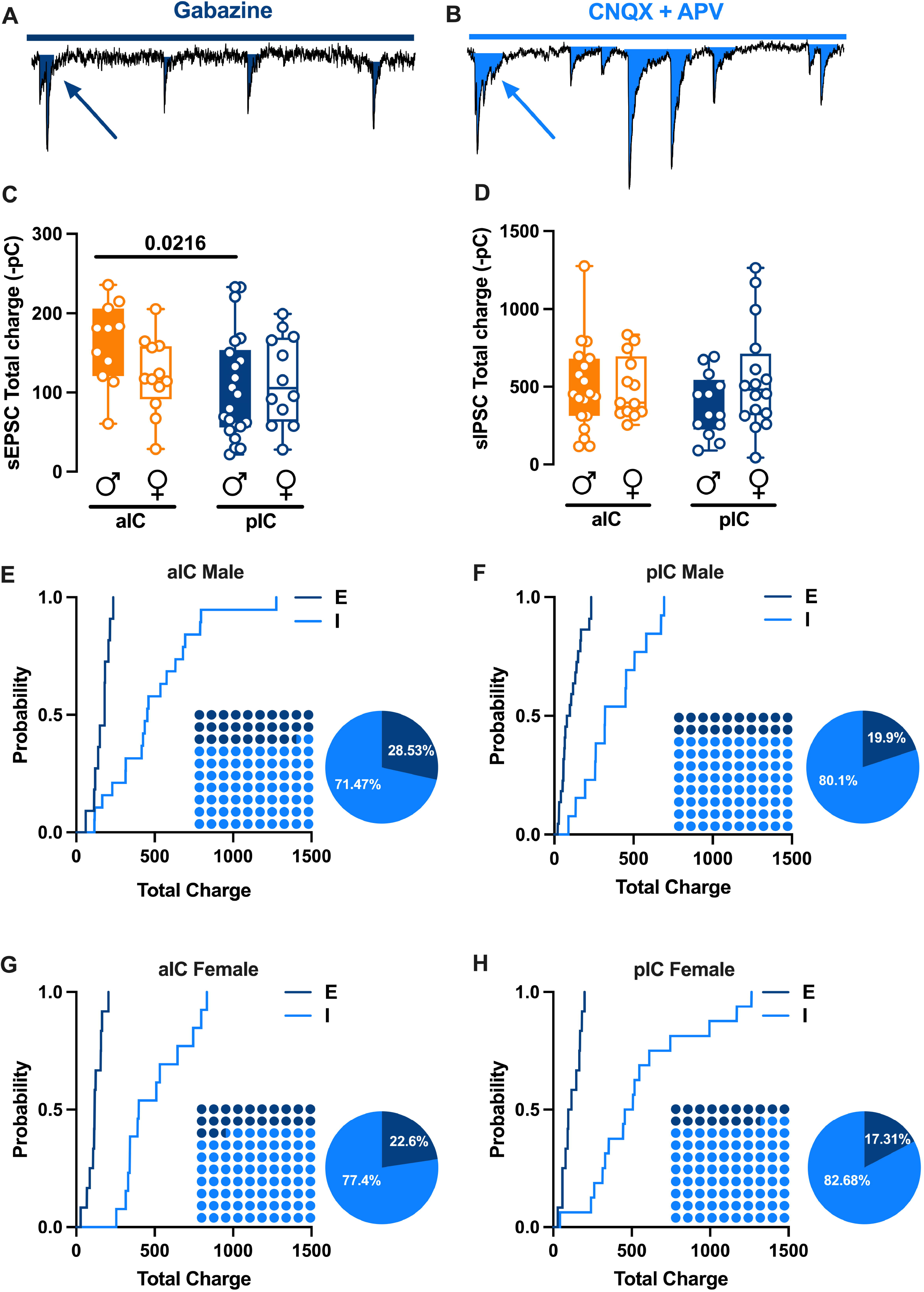
Sex and subregional profiles of the Excitatory/Inhibitory (E/I) balance of the Insular Cortex. **(A-B)** Schematic illustration of synaptic charge tranfered by each sEPSCs (A) and sIPSCs (B) calculated as the area inside each event as indicated by the arrows. **(C)** An analysis of the total charge of AMPA-sEPSCs measured over a 6-minute period across sexes revealed similar total charges in both male and female in both the aIC and pIC. However, there was a higher total charge transfer observed in the aIC compared with pIC, but this was only observed in males. **(D)** Similarly, the total charge of GABA-sIPSCs, when measured over a 6-minute period considering both area and sex, showed a comparable amount of charge transferred. This was analogous to the observations made in **’C’** for AMPA-sEPSCs. **(E-H)** Cumulative frequency distribution of sEPSCs (E) and sIPSCs (I) total charge transfer obtain from each Insular cortex within male and female animals. Insets: dot plots and pie graph showing the proportion of E versus I extrapolated at *P* = 0.5 from the corresponding cumulative frequency. **(C-D)** Data are presented as box-and-whisker plots (minimum, maximum, median) and analyzed via two-way ANOVA followed by Šídák’s multiple comparison. P-values <0.05 are depicted in the graphs. **(C)** The sample sizes for aIC male were 12/10 for pIC male 22/15, for aIC female 13/10 and for pIC female 12/10. **(D)** The sample sizes for aIC male were 19/11 for pIC male 13/9, for aIC female 13/10 and for pIC female 16/10.

**Table 4.** Total charge transferred by AMPA- and GABA-mediated events in aIC and pIC of both sexes.

| Misure | Area | Sex | Median | MAX | MIN | n/N | Two-way<br>Anova<br>Main effects<br>and<br>interactions | Multiple<br>comparison<br>(Šidák's post<br>hoc test) |
| --- | --- | --- | --- | --- | --- | --- | --- | --- |
| sEPSCs<br>Total<br>charge<br>(-pC) | aIC | M | -180894 | -60757 | -235773 | 12/10 | Interaction:<br>p = 0.1101<br>Area:<br>F (1. 53) =<br>3.680<br>p = 0.06<br>Sex:<br>F (1. 53) =<br>1.017<br>p = 0.3178 | aIC Male vs pIC<br>Male<br>p = 0.0216 |
|  |  | F | -117082 | -28617 | -205287 | 13/10 |  | aIC Female vs<br>pIC Female<br>p = 0.9756 |
|  | pIC | M | -88857 | -21648 | -233047 | 22/15 |  | aIC Male vs aIC<br>Female<br>p = 0.1688 |
|  |  | F | -105611 | -27801 | -199381 | 12/10 |  | pIC Male vs pIC<br>Female<br>p = 0.8692 |
| sIPSCs<br>Total<br>charge<br>(-pC) | aIC | M | -453628 | -115748 | -1275617 | 19/11 | Interaction: p =<br>0.2221<br>Area:<br>F (1. 57) =<br>0.1999 p =<br>0.6565<br>Sex:<br>F (1. 57) =<br>1.486 p =<br>0.2278 | aIC Male vs pIC<br>Male<br>p = >0.9999 |
|  |  | F | -398812 | -255849 | -833222 | 13/10 |  | aIC Female vs<br>pIC Female<br>p = 0.9756 |
|  | pIC | M | -319953 | -89489 | -692799 | 13/9 |  | aIC Male vs aIC<br>Female<br>p = 0.4082<br>pIC Male vs pIC<br>Female<br>p = 0.1789 |

### Subregional differences of excitatory synaptic transmission

To explore the hypothesis that synaptic connectivity is responsible for the observed subregional differences in E/I balance, we conducted recordings and compared spontaneous AMPA-mediated post-synaptic currents in layer V pyramidal neurons within both sexes and subregions (**Fig. 5-6**). In males, it became evident that the mean amplitude of post-synaptic currents in the pIC increased, while the mean frequency was lower compared to the anterior aIC (**Fig. 5C-E**, **Table 5**). Further analysis of the frequency distribution confirmed a higher proportion of larger but less frequent excitatory events in pIC compared to aIC (**Fig. 5D-F**). In contrast, for females, the mean amplitude and frequency of spontaneous excitatory post-synaptic currents (sEPSCs) remained similar in both aIC and pIC (**Fig. 6C-E**, **Table 5**). A closer examination of the frequency distribution of individual sEPSCs, however, revealed a higher proportion of smaller and more frequent synaptic currents in aIC compared to pIC in females (**Fig. 6D-F, Table5**). Notably, when comparing across sexes and subregions, a sex-specific difference was observed solely in pIC. Specifically, the primary frequency of sEPSCs was higher in females than in males (**Supplementary Fig. 2B, Suppl. Table 2**). Finally, no significant differences were found in the kinetics of excitatory events across both subregions and sexes (**Supplementary Fig. 2C-D, Suppl. Table 2**)

**Figure 5:**
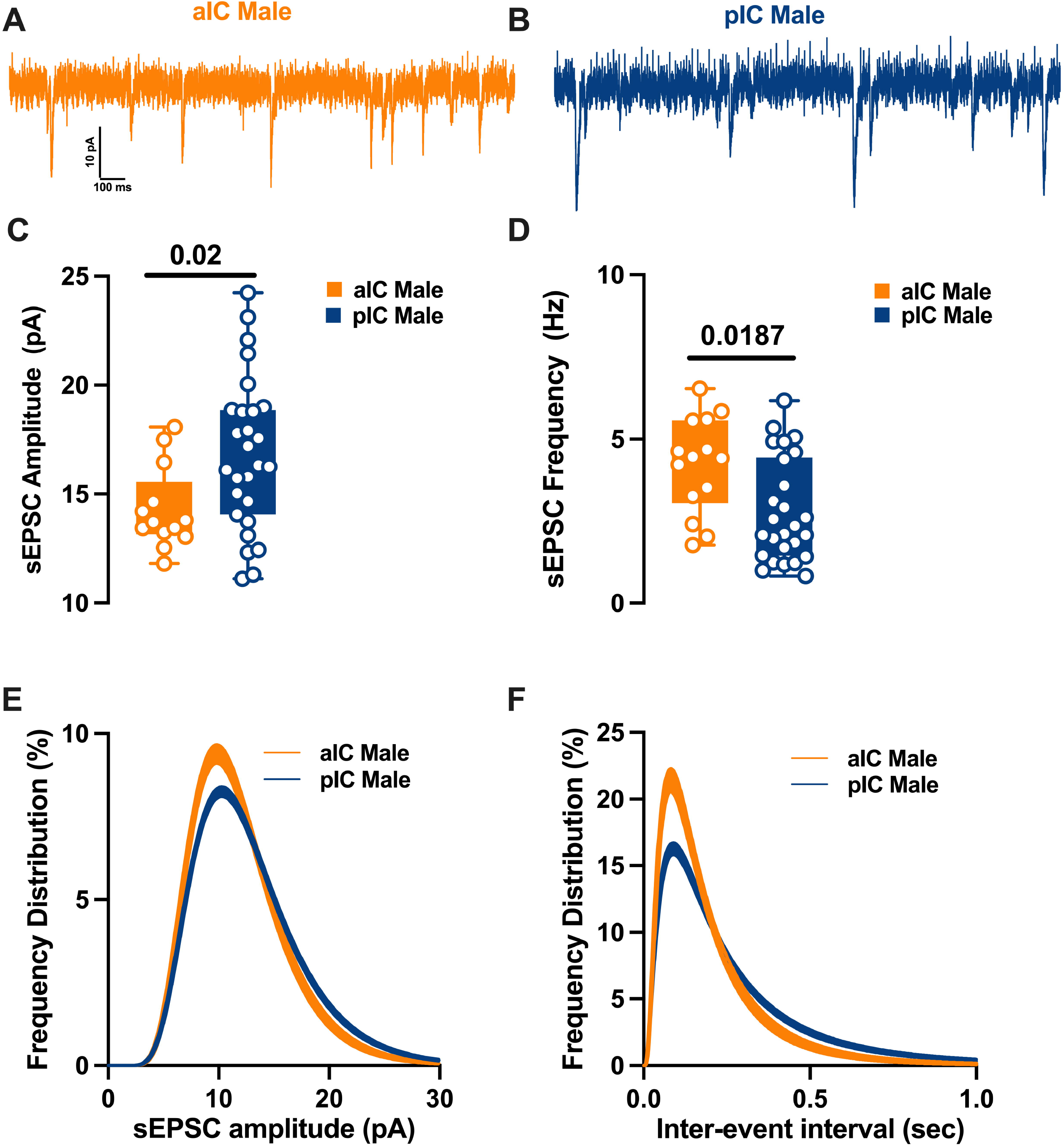
Region-specific spontaneous excitatory synaptic activity in adult male IC pyramidal neurons. **(A-B)** Representative spontaneous excitatory postsynaptic currents (sEPSCs) recorded at -70 mV in male aIC and pIC. **(C-D)** On average, excitatory events are larger and less frequent in the pIC than the aIC. **(E-F)** Log-normal curve fittings with confidence intervals (± CI) reveal that the amplitude and frequency distribution are skewed to the right in the pIC as compared to the aIC. Data are presented as box-and-whisker plots (minimum, maximum, median) and analyzed via Mann-Whitney U test. P-values <0.05 are depicted in the graphs. The sample size for aIC male was 14/9, and for pIC male was 27/16.

**Figure 6:**
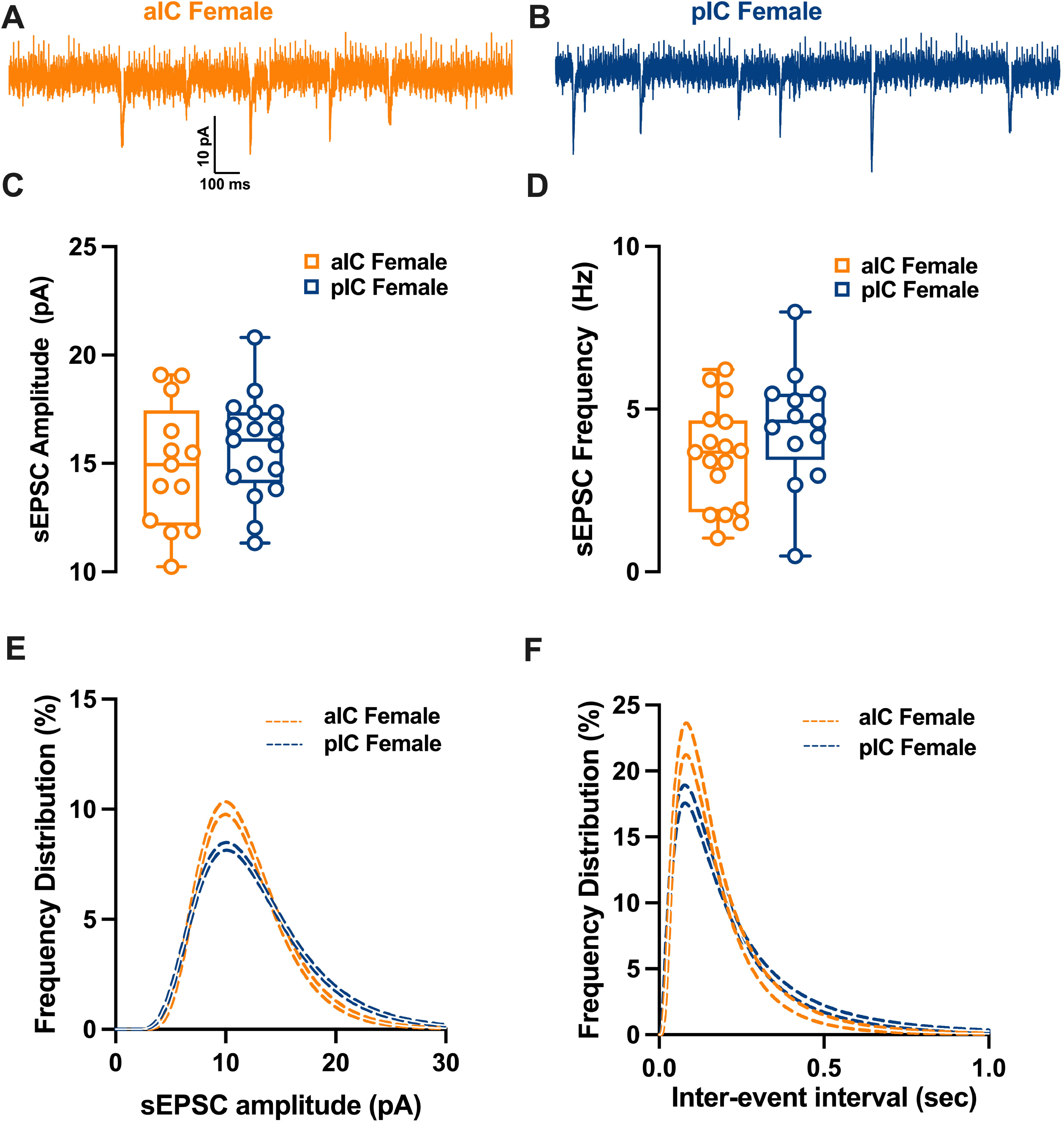
Differences in excitatory synaptic activity in female insular pyramidal neurons by region. **(A-B)** Representative spontaneous excitatory postsynaptic currents (sEPSCs) recorded at -70 mV in female aIC and pIC. **(C-D)** On average, excitatory events are comparable in amplitude and frequency in both aIC and pIC. **(E-F)** Log-normal curve fittings with confidence intervals (± CI) reveal higher proportion of smaller and more frequent events in the aIC as compared to the pIC. Data are presented as box-and-whisker plots (minimum, maximum, median) and analyzed via Mann-Whitney U test. P-values <0.05 are depicted in the graphs. The sample size for aIC female was 13/6, and for pIC female was 17/12.

**Table 5.** sE/IPSCs within the IC in both sexes.

| Misure | Sex | aIC |  |  |  | pIC |  |  |  | p value<br>Multiple<br>Mann-Whitney |
| --- | --- | --- | --- | --- | --- | --- | --- | --- | --- | --- |
|  |  | Median | MAX | MIN | n/N | Median | MAX | MIN | n/N |  |
| sEPSCs Amplitude (pA) | M | 13.71 | 18.09 | 11.82 | 14/9 | 16.33 | 24.25 | 11.12 | 27/16 | 0,0229 |
| sEPSCs Frequency (Hz) | M | 4.4 | 6.53 | 1.77 |  | 2.235 | 6.17 | 0.83 |  | 0,0116 |
| sEPSCs Amplitude (pA) | F | 14.95 | 19.09 | 10.24 | 13/6 | 16.08 | 20.81 | 11.33 | 17/12 | 0,3851 |
| sEPSCs Frequency (Hz) | F | 3.68 | 6.22 | 1.04 |  | 4.63 | 7.99 | 0.49 |  | 0,1252 |
| sIPSCs Amplitude (pA) | M | 38.83 | 68.11 | 18.89 | 22/10 | 35.91 | 62.49 | 18.36 | 14/11 | 0,4507 |
| sIPSCs Frequency (Hz) | M | 5.17 | 8.32 | 1.75 |  | 2.93 | 6.54 | 1 |  | 0,0418 |
| sIPSCs Amplitude (pA) | F | 44.53 | 64.1 | 23.39 | 15/10 | 34.58 | 67 | 18.89 | 19/11 | 0,0876 |
| sIPSCs Frequency (Hz) | F | 4.12 | 8.05 | 1.88 |  | 4.39 | 10.08 | 0.62 |  | >0,9999 |

### Differential GABAergic Transmission in aIC and pIC of Adult Males

Spontaneous GABA_A_-mediated inhibitory post-synaptic currents were recorded in IC principal neurons in whole-cell configuration (**Fig. 7-8**). As shown in **Fig.7C**, in adult male the mean amplitude of sIPSCs remained similar among the two cortexes. In contrast the frequency was lower in pIC when compared to those in aIC (**Fig. 7E**, **Table 5**). Indeed, the frequency distribution confirmed that inhibitory events were more frequent in aIC (**Fig. 7F**). In contrast, in female group both mean amplitude and frequency were comparable (**Fig. 7C-E**, **Table 5**). When compared across sex and subregion no further differences were found in the mean amplitude, frequency or kinetics of sIPSCs (**Supplementary. Fig. 3A-D, Suppl. Table 2**).

**Figure 7:**
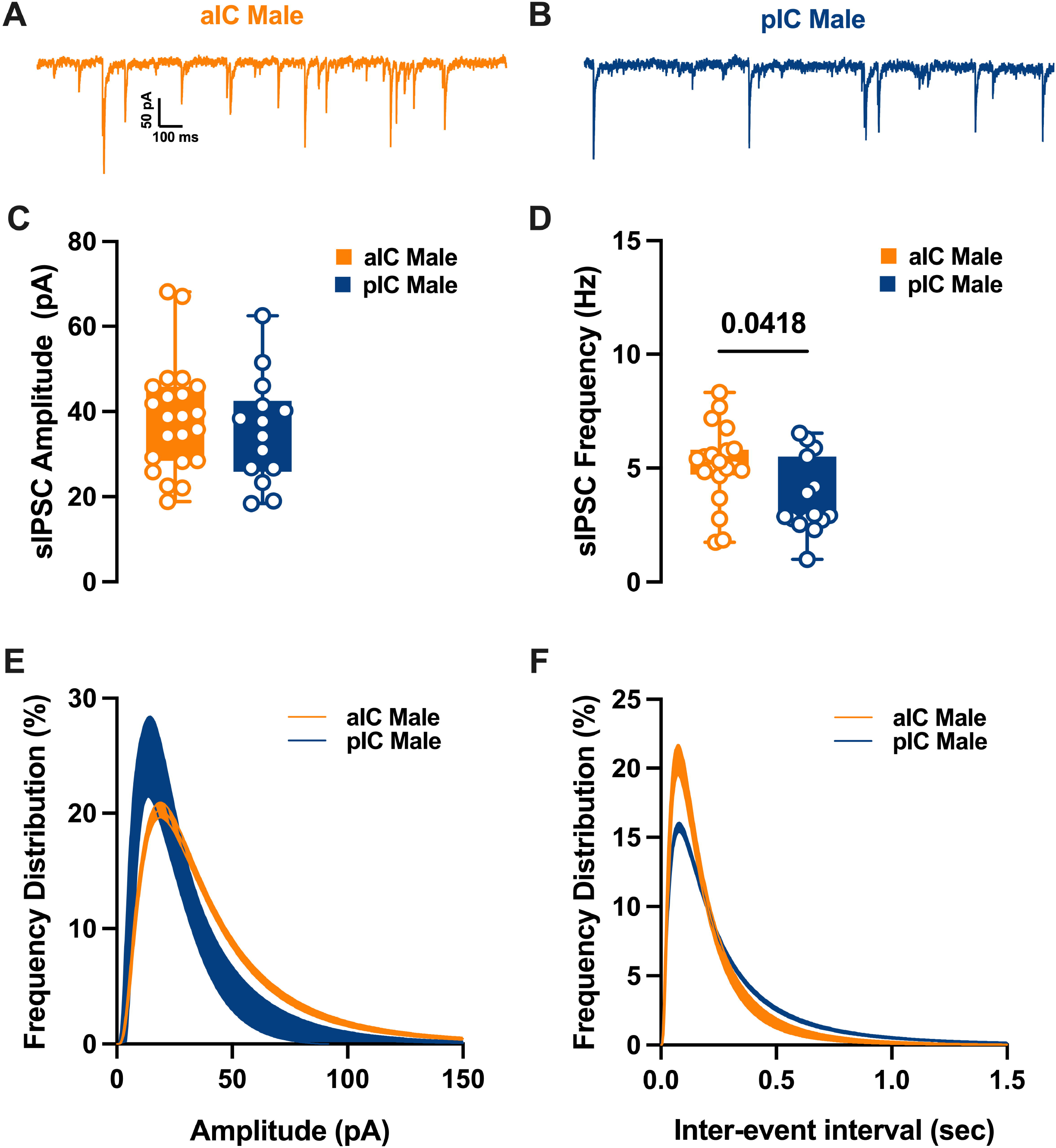
Differences in inhibitory synaptic activity among male pyramidal neurons in the insular cortex, by region. **(A-B)** Representative spontaneous inhibitory postsynaptic currents (sIPSCs) recorded at -70 mV in male aIC and pIC. **(C-D)** Upon conducting a quantitative analysis of the mean amplitude and frequency across these areas, it was found that although the amplitudes were similar, there was a lower frequency of inhibitory events in the pIC when compared to the aIC. **(E-F)** Log-normal curve fittings with confidence intervals (± CI) reveal that the amplitude and frequency distribution are skewed to the right and to the left respectively in the aIC as compared to the pIC. Data are presented as box-and-whisker plots (minimum, maximum, median) and analyzed via Mann-Whitney U test. P-values <0.05 are depicted in the graphs. The sample size for aIC male was 22/10, and for pIC male was 14/11.

**Figure 8:**
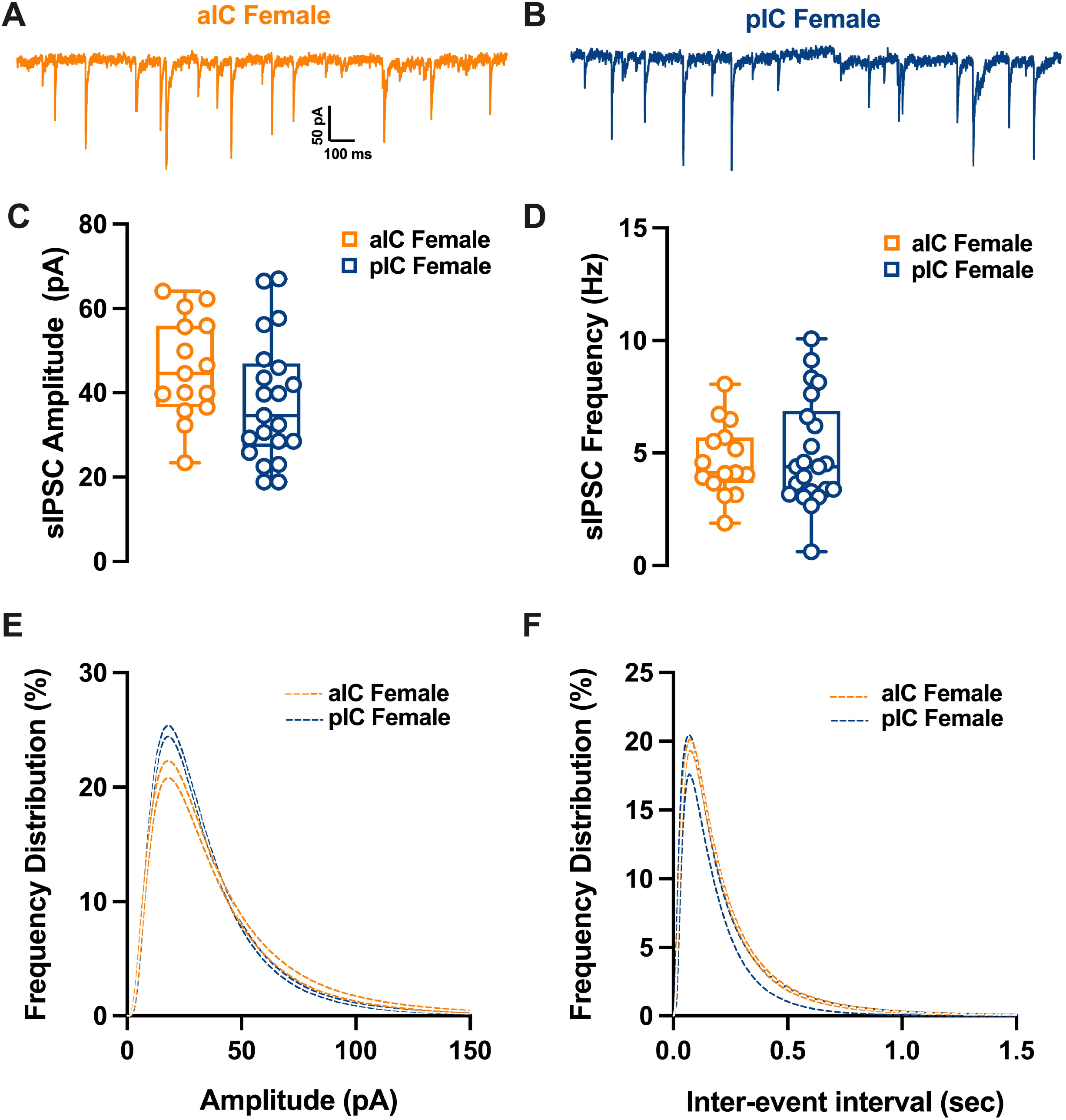
Similarities in inhibitory synaptic activity among female pyramidal neurons across different subregions of the insular cortex. **(A-B)** Representative spontaneous inhibitory postsynaptic currents (sIPSCs) recorded at -70 mV in female aIC and pIC. **(C-D)** On average, ihibitory events are comparable in amplitude and frequency in both aIC and pIC. **(E-F)** Log-normal curve fittings with confidence intervals (± CI) reveal higher proportion of smaller events in pIC compared to aIC. Data are presented as box-and-whisker plots (minimum, maximum, median) and analyzed via Mann-Whitney U test. P-values <0.05 are depicted in the graphs. The sample size for aIC female was 15/10, and for pIC female was 19/11.

### LTP at the glutamatergic synapses in a sex- and subregion-specific manner

The IC exhibits reciprocal connections with cortical and subcortical regions associated with sensory, cognitive, and memory functions, suggesting the involvement of synaptic plasticity mechanisms. While Long-Term Potentiation (LTP) has been studied in the aIC, our understanding of LTP in the pIC is limited. Additionally, there is a paucity of knowledge regarding potential sex differences in synaptic processes within the IC. Thus, excitatory post synaptic transmission in each IC were recorded in layer V pyramidal neuron in both male and female (**Fig. 9A-B**). We fist compared input-output (I/O) curves of fEPSP (**Fig. 9C-D**, **Table 6**). The I/O relationship did not differ between aIC and pIC in male (**Fig. 9C**), in contrast female showed a marked difference across subregions. Indeed, along the entire I/O curve the aIC showed a greater synaptic strength compared to the pIC (**Fig. 9D**). LTP could be elicited in both aIC and pIC in male (**Fig. 9E-F**) and in contrast, LTP could only be induced in the aIC of female (**Fig. 9G-H**).

**Figure 9:**
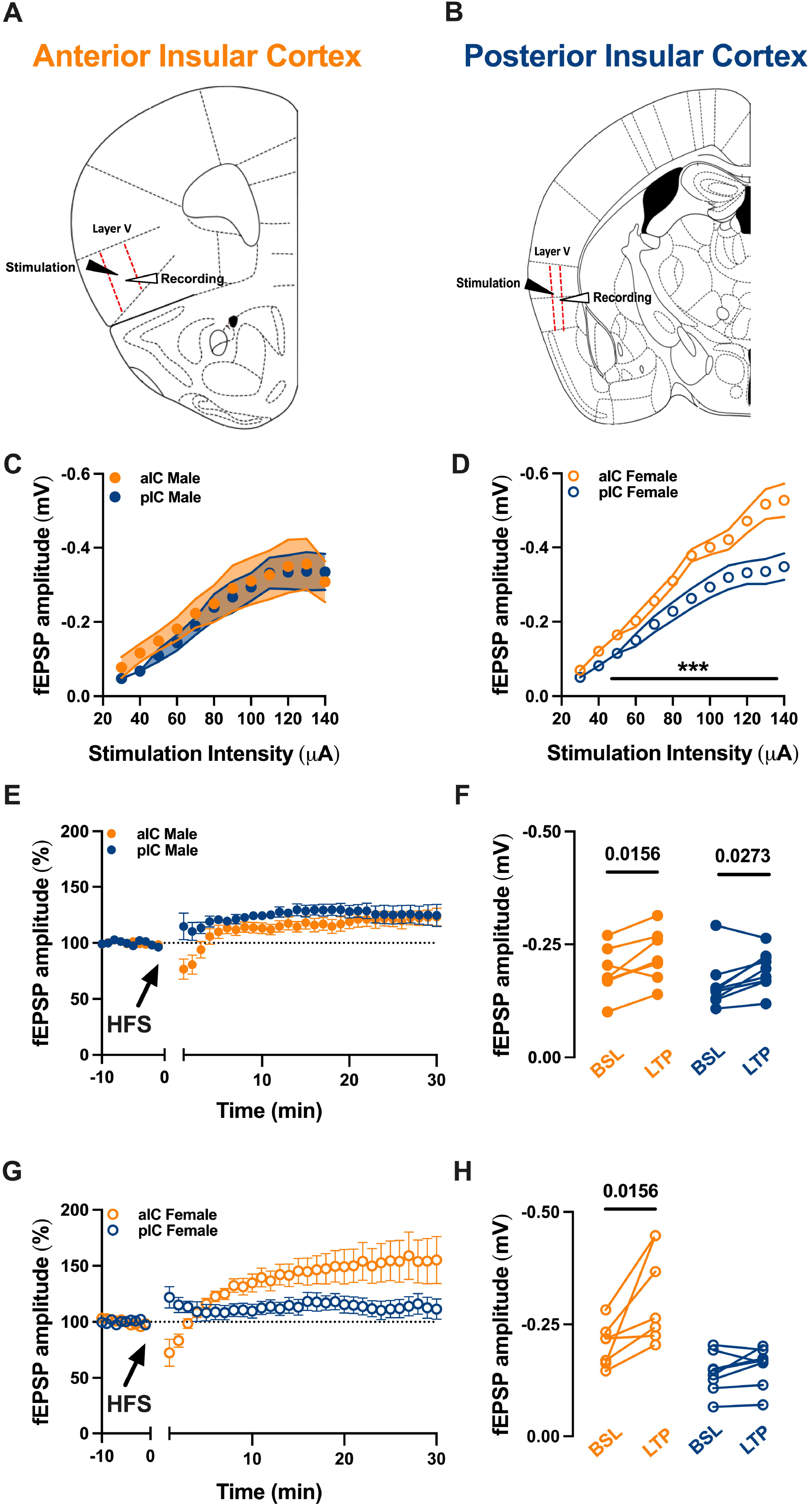
Long-term potentiation in the insular cortex varies based on sex and subregion. **(A-B)** A schematic illustration indicates the aIC and pIC, both delineated by red dashed lines. Stimulation and recordings were applied and gathered respectively from layer V pyramidal neurons in each insular cortex. **(C-D)** An examination of the input-output (I/O) relationship showed a comparable synaptic strength in both the aIC and pIC of adult males **(C)**. Conversely, female aIC exhibited a greater synaptic strength in comparison to pIC. **(E-G)** The field excitatory postsynaptic potential (fEPSP), shown as a percentage of the baseline, was observed before and after the application of high-frequency stimulation (HFS). The point in time at which HFS was applied is shown by the arrow. **(E)** When HFS protocol was administered to layer V pyramidal neurons in adult males, it led to long-term potentiation (LTP) in both the anterior and posterior insular cortices. **(F)** The fEPSP magnitude at baseline (from 10 to 0 minutes) and during LTP (from 20 to 30 minutes after induction), corresponding to the normalized values in ’C’, demonstrated a notable difference in the 10-minute baseline period and the final 10 minutes of recording in both male insular cortices. **(G)** In contrast, the same protocol, when applied in adult females, only induced a strong LTP in the aIC. **(H)** The fEPSP magnitude at baseline (from 10 to 0 minutes) and during LTP (from 20 to 30 minutes post-induction), corresponding to the normalized values in ’F’, showed a significant difference in the 10-minute baseline period and the final 10 minutes of recording, but only in the aIC of adult females. (**C-D-E-G**) Data are presented as mean ± SEM in XY plots and analyzed via Mann-Whitney U test (**C-D**). P-values <0.05 are depicted in the graphs. **(F-H)** Data are shown as pre-post individual experiments and analyzed via Wilcoxon P test. P-values <0.05 are depicted in the graphs. The sample sizes for aIC and pIC male and female were 8 and 7, respectively.

**Table 6.** Input-output (I/O) Relationship of aIC and pIC of adult males and females.

| Measure<br>(unit) | Condition | Injected<br>current<br>(μA) | aIC | | | pIC | | | *p value*<br>*Multiple*<br>*Mann-*<br>*Whitney*<br>*Unpaired t*<br>*test* |
| --- | --- | --- | --- | --- | --- | --- | --- | --- | --- |
|  |  |  | Mean | SEM | n | Mean | SEM | n |  |
| **I/O** | **Male** | 30 | -0.07 | 0.02 | 7 | -0.04 | 0.004 | 8 | 0.53 |
|  |  | 40 | -0.11 | 0.024 |  | -0.06 | 0.009 |  | 0.12 |
|  |  | 50 | -0.14 | 0.02 |  | -0.10 | 0.015 |  | 0.24 |
|  |  | 60 | -0.18 | 0.03 |  | -0.14 | 0.02 |  | 0.32 |
|  |  | 70 | -0.22 | 0.03 |  | -0.18 | 0.022 |  | 0.66 |
|  |  | 80 | -0.24 | 0.04 |  | -0.23 | 0.039 |  | >0.999999 |
|  |  | 90 | -0.29 | 0.06 |  | -0.26 | 0.04 |  | >0.999999 |
|  |  | 100 | -0.31 | 0.06 |  | -0.29 | 0.03 |  | 0.79 |
|  |  | 110 | -0.32 | 0.06 |  | -0.33 | 0.04 |  | 0.79 |
|  |  | 120 | -0.34 | 0.07 |  | -0.33 | 0.04 |  | 0.79 |
|  |  | 130 | -0.35 | 0.06 |  | -0.33 | 0.05 |  | 0.93 |
|  |  | 140 | -0.30 | 0.05 |  | -0.33 | 0.047 |  | 0.69 |

## Discussion

The principal findings of this study reveal distinct electrophysiological characteristics between the aIC and pIC in both males and females. These differences encompass the properties of pyramidal neurons, with aIC neurons being larger and less excitable than those in pIC. Our study highlights that the E/I balance within the IC leans towards inhibition, with pIC exhibiting a higher degree of inhibitory activity compared to aIC, regardless of sex. Finally, there are sex-specific distinctions in synaptic plasticity.

Examination of the passive and active properties of layer V (L5) pyramidal neurons, revealed that in both sexes, principal neurons exhibit distinct characteristics along the rostro-caudal axis. Specifically, pyramidal neurons in the aIC are bigger and more hyperpolarized compared to those in the posterior pIC. Consequently, due to their different resting potentials, pyramidal neurons in aIC display a higher rheobase compared to those in pIC. Although no differences were observed in the voltage membrane response between aIC and pIC in males, adult females exhibited a lower membrane response in aIC compared to pIC. These subregional variances in IC pyramidal neurons can be attributed to the distinct morphological and physiological properties associated with cortical neurons projecting to various cortical and subcortical targets ^14^. Firing patterns primarily depend on a cell’s intrinsic properties ^15–17^. To investigate whether the differences in passive and active membrane properties between aIC and pIC influence the intrinsic excitability of L5 pyramidal neurons in both IC subregions, we compared their excitability. In line with the more hyperpolarized resting potential and higher rheobase, pyramidal neurons in aIC exhibited lower excitability compared to those in pIC, regardless of sex.

The IC follows a posterior-to-anterior gradient, with pIC being highly excitable and aIC being less excitable. Given the strong association between the functional output of a brain region and the intrinsic excitability of its neurons, it’s reasonable to propose that the heightened excitability observed in pIC could be linked to its function as the primary input hub for sensory signals originating from the sensory cortex and sensory thalamic nuclei ^10^. The excitatory and E/I balance is a crucial factor in brain function, yet the E/I balance within the IC had not been previously characterized. Our investigation uncovered a prevalence of inhibitory signals over excitatory ones in both the aIC and pIC. Notably, pIC exhibited a higher proportion of inhibitory activity in both males and females, suggesting that inhibitory processes may play a more significant role in this subregion. This inhibitory tone is believed to function as a filter for synchronizing excitatory activity, enhancing temporal precision and the dynamic range of firing to coordinate responses to sensory input ^18,19^. Our findings that the E/I balance within the IC tends to lean towards inhibition is consistent with previous observations in conditions related to the IC, such as pain-related disorders ^11^. The data also revealed a sex-specific difference in the I/O relationship. While males exhibited comparable synaptic strength and LTP, females displayed greater synaptic strength in aIC compared to pIC. In females, the high-frequency stimulation protocol led to LTP in aIC only. These differences in synaptic plasticity may underlie distinct processes involved in multisensory integration in males and females ^20–23^.

### Perspectives and Significance

This research provides invaluable insights into the intricate role of the IC, a central hub for processing emotional and interoceptive information, which holds direct relevance in comprehending sex-based disparities in behavior and cognition. Through the examination of distinctive properties of pyramidal neurons within both the anterior and posterior IC, our study not only underscores sex-specific variations but also unveils their potential implications on intrinsic properties, synaptic plasticity, and excitatory transmission. These findings offer a fresh perspective on the neural underpinnings of sex-related differences in behavior and cognition, holding the promise of extending their impact to the development of more precisely targeted interventions and treatments. This is particularly pertinent in disorders where the IC plays a pivotal role, given that these conditions often exhibit a pronounced sex bias, including prevalent instances such as anxiety, depression, and somatosensory processing disorders.

## Acknowledgements

The authors are grateful to the Chavis-Manzoni team members for helpful discussions.

## Authors’ contributions

DI: Conceptualization, Data curation, Formal analysis, Validation, Writing—original draft, review and editing. AC: Formal analysis. BS: Statistical analysis. PC: Conceptualization, Supervision. OJM: Conceptualization, Supervision, Funding acquisition, Methodology, Project administration, Writing—original draft, review, and editing.

## Ethics declarations

### Ethics approval and consent to participate

Animals were treated in compliance with the European Communities Council Directive (86/609/EEC) and the United States National Institutes of Health Guide for the care and use of laboratory animals. All procedures using experimental animals were approved by Aix-Marseille University Institutional Animal Care & Use Committee.

### Consent for publication

All authors read and approved the final manuscript for publication.

### Availability of data and materials

All data reported in this paper will be shared by the lead contact upon request. This paper does not report original code. Any additional information required to reanalyze the data reported in this paper is available from the lead contact upon request.

## Supplementary figure legends

**Supplemental Figure 1: Comparative analysis of I-V relationships and excitability in male and female anterior and posterior IC neurons. (A-B**) Incremental current injections of 50 pA from -400 pA to +50 pA exhibited no significant differences in the I-V relationship for male and female neurons in both anterior and posterior IC. **(C-D**) Gradual depolarizing current injections (500 ms, ranging from 0 pA to 600 pA in 50 pA increments) revealed comparable excitability in both male and female neurons within the same area. **(A-B)** Each dot similarly represents the group mean value at the given current step, with data shown as mean ± SEM in an XY plot. A Mann-Whitney U test was used, and *p-value <0.05 was considered significant. **(C-D**) Each dot indicates the group mean value for the respective current step, with data presented as mean ± CI in an XY plot. A Mann-Whitney U test was used for statistical analysis, with a *p-value <0.05 considered significant. **(A-D)** aIC male is represented as 10/14 in dark orange, aIC female as 6/14 in light orange, pIC male as 15/20 in dark blue, and pIC female as 12/16 in light blue.

**Supplemental Figure 2: Quantitative evaluation of amplitude, frequency, and kinetics of excitatory events in adult male and female anterior and posterior IC neurons.** A quantitative analysis of mean amplitude and frequency in relation to area and sex demonstrated greater amplitude **(A)** but lower frequency **(B)** of pIC excitatory events compared to aIC events in adult males only. **(B)** In females, pIC was characterized by higher excitatory transmission compared to male pIC. **(C-D)** The kinetics of sEPSC were consistent across both areas and sexes. **(A-D**) Individual neurons are represented by single dots. Data are displayed as box and whisker plots (min., max., median). A two-way ANOVA followed by a Šídák multiple comparison test was used for data analysis. P-values <0.05 are indicated in the graphs. aIC male is represented as 9/14 in dark orange, pIC male as 16/27 in dark blue, aIC female as 6/13 in light orange, and pIC female as 12/17 in light blue.

**Supplemental Figure 3: Quantitative assessment of sIPSC events in male and female anterior and posterior IC neurons: inhibitory transmission comparison.** A quantitative analysis of sIPSC events in relation to area and sex showed similar amplitude of sICPCs in both males and females **(A),** while there was a lower frequency of inhibitory events in the pIC when compared to the aIC, in only male **(B)**. No differences were found in the kinetic of sIPSCs **(C-D)**. Each dot represents an individual neuron. Data are displayed as box and whisker plots (min., max., median). A two-way ANOVA followed by a Šídák multiple comparison test was used for data analysis. P-values <0.05 are indicated in the graphs. aIC male is represented as 10/21 in dark orange, pIC male as 11/15 in dark blue, aIC female as 10/16 in light orange, and pIC female as 11/19 in light blue.

