## Supplementary figures and images for "Sexual differences in neuronal and synaptic properties across subregions of the mouse insular cortex"

### sup fig 1

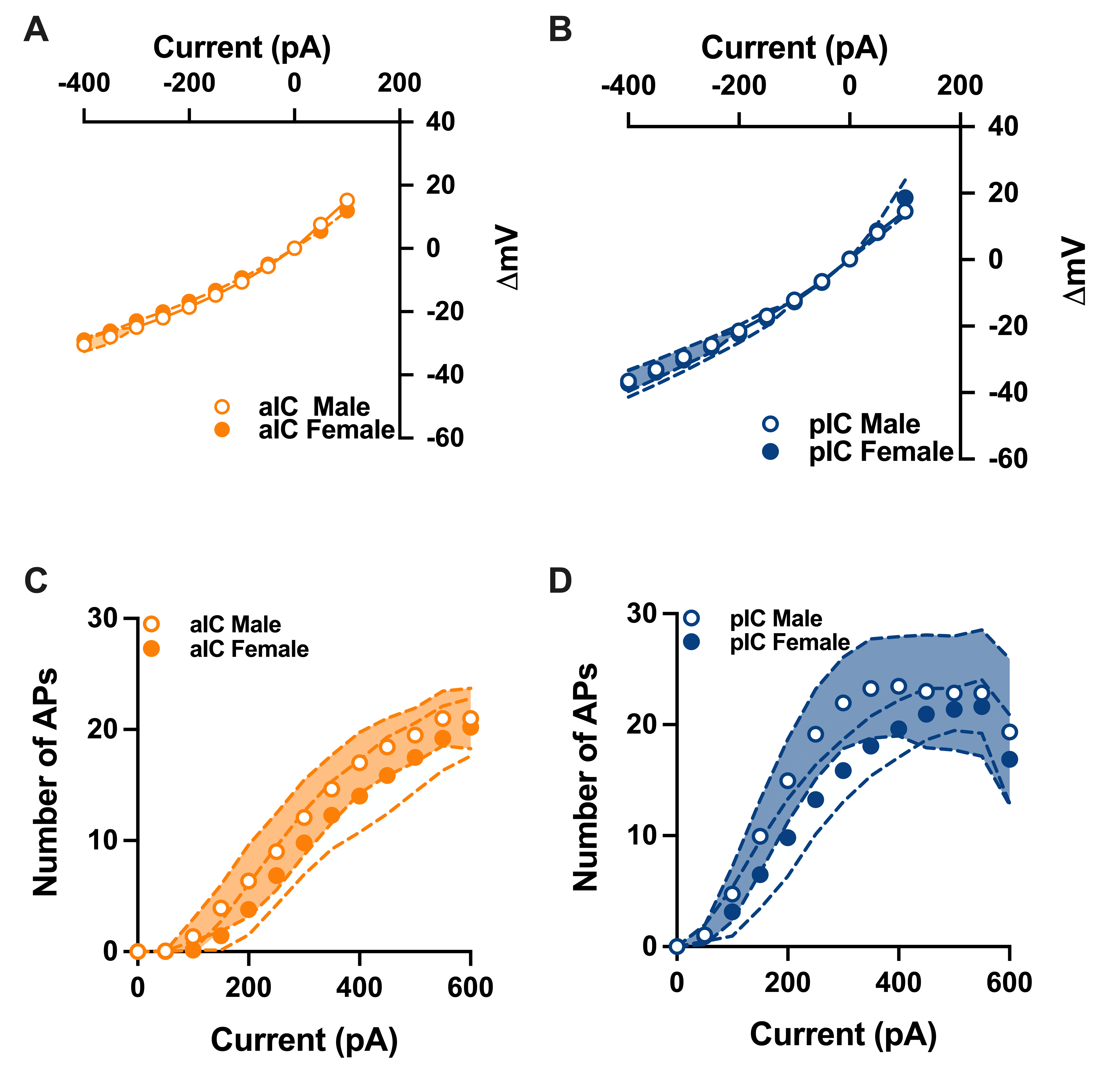

### sup fig 2

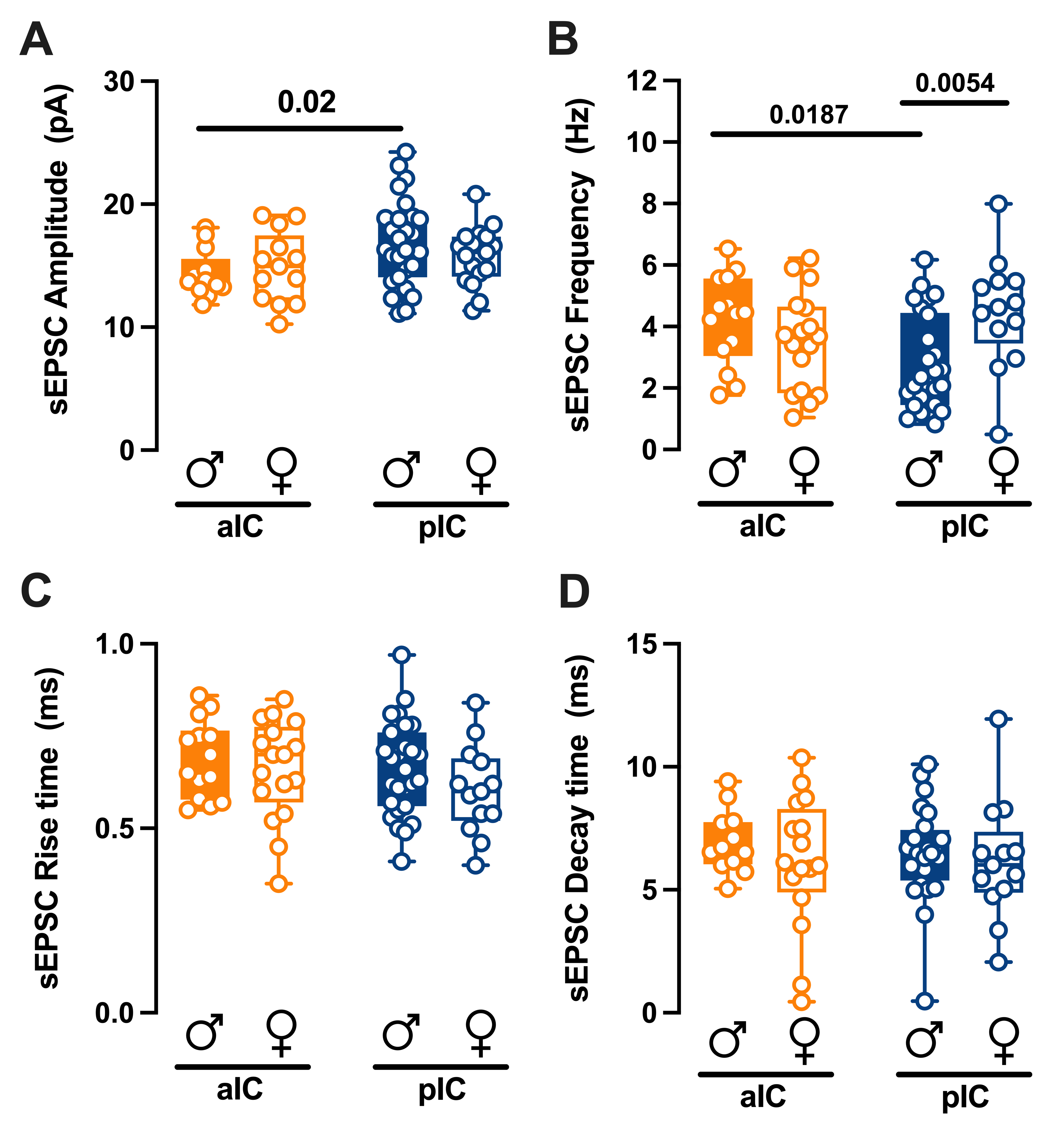

### sup fig 3

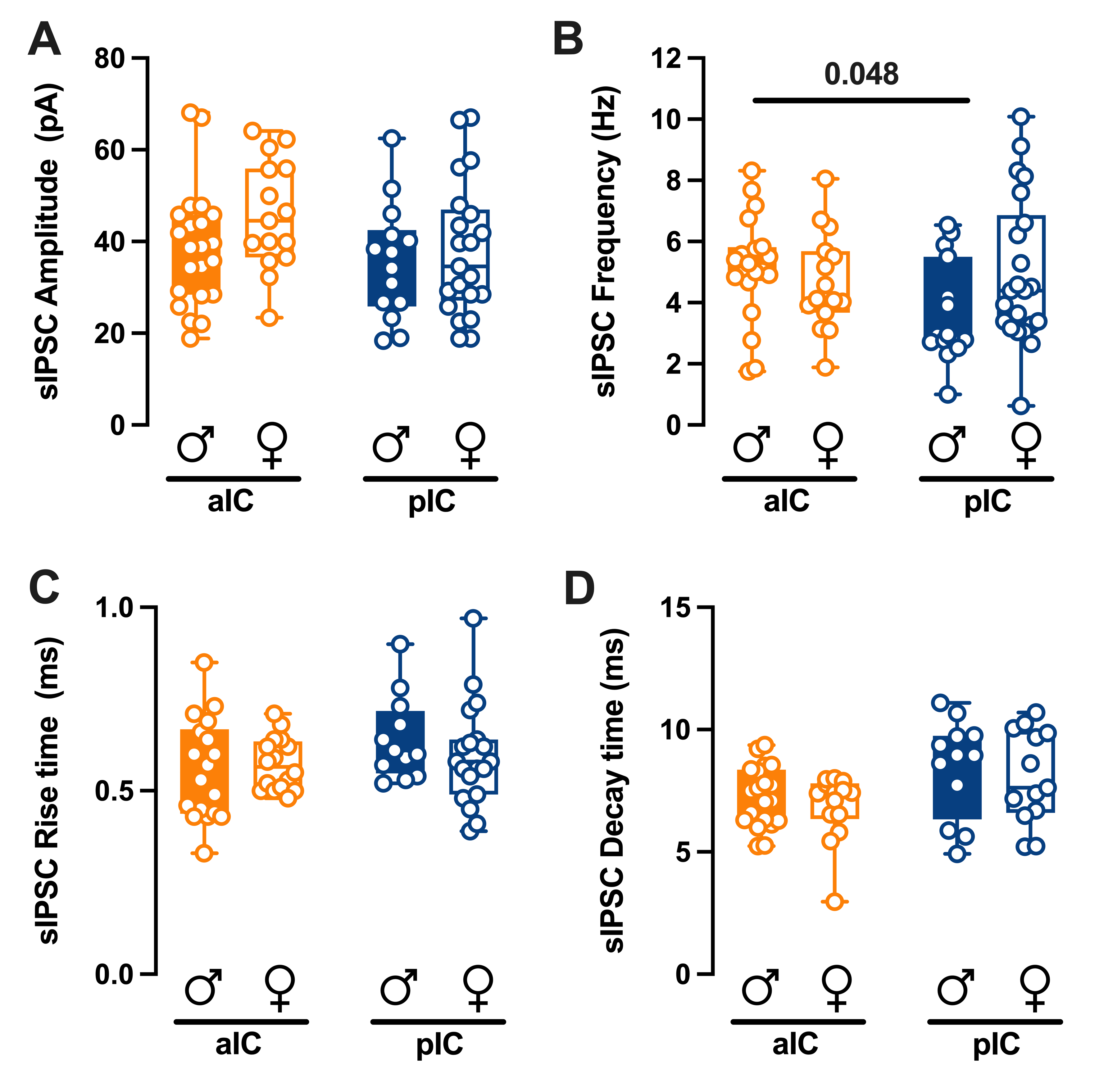
