## Supplementary material for "Sexual differences in neuronal and synaptic properties across subregions of the mouse insular cortex": sup table 1

**Supplementary Table 1. Comparative analysis of I-V relationships and excitability in male and female anterior and posterior IC neurons.**

| **Measure**  (unit) | **Area** | **Injected  current** (pA) | **MALE** | | | **FEMALE** | | | ***p value  Multiple Mann-Whitney  Unpaired t test*** |
| --- | --- | --- | --- | --- | --- | --- | --- | --- | --- |
|  |  |  | Mean | SEM | n/N | Mean | SEM | n/N |  |
| **Voltage changes per current step (ΔVm)** | **aIC** | *-400* | -30.53 | 2.28 | 14/10 | -28.93 | 1.65 | 13/6 | 0.54 |
|  |  | *-350* | -27.97 | 2.18 |  | -26.19 | 1.6 |  | 0.43 |
|  |  | *-300* | -24.88 | 1.97 |  | -22.97 | 1.21 |  | 0.48 |
|  |  | *-250* | -21.97 | 1.79 |  | -20.08 | 1.2 |  | 0.43 |
|  |  | *-200* | -18.51 | 1.59 |  | -16.75 | 1.1 |  | 0.51 |
|  |  | *-150* | -14.76 | 1.31 |  | -13.31 | 0.86 |  | 0.57 |
|  |  | *-100* | -10.61 | 1 |  | -9.3 | 0.62 |  | 0.51 |
|  |  | *-50* | -5.67 | 0.58 |  | -5.01 | 0.38 |  | 0.57 |
|  |  | *0* | 0.047 | 0.04 |  | 0.09 | 0.08 |  | 0.98 |
|  |  | *50* | 7.61 | 1.21 |  | 5.52 | 0.47 |  | 0.31 |
|  |  | *100* | 15.21 | 1.68 |  | 11.92 | 0.99 |  | 0.31 |
|  | **pIC** | *-400* | -36.49 | 3.14 | 28/15 | -37.23 | 1.94 | 20/12 | 0.31 |
|  |  | *-350* | -32.99 | 2.82 |  | -33.91 | 1.78 |  | 0.25 |
|  |  | *-300* | -29.29 | 2.50 |  | -30.23 | 1.61 |  | 0.28 |
|  |  | *-250* | -25.65 | 2.21 |  | -26.34 | 1.43 |  | 0.34 |
|  |  | *-200* | -21.45 | 1.84 |  | -22.33 | 1.28 |  | 0.35 |
|  |  | *-150* | -16.94 | 1.50 |  | -17.74 | 1.1 |  | 0.3 |
|  |  | *-100* | -12.14 | 1.13 |  | -12.68 | 0.86 |  | 0.39 |
|  |  | *-50* | -6.49 | 0.62 |  | -6.85 | 0.55 |  | 0.43 |
|  |  | *0* | 0.15 | 0.06 |  | 0.21 | 0.15 |  | 0.89 |
|  |  | *50* | 8.12 | 0.92 |  | 8.74 | 0.97 |  | 0.61 |
|  |  | *100* | 14.57 | 1.18 |  | 18.63 | 2.57 |  | 0.43 |
| **Number of APs** | **aIC** | *0* | 0 | 0 | 14/10 | 0 | 0 | 14/6 | >0.999999 |
|  |  | *50* | 0.07 | 0.07 |  | 0 | 0 |  | >0.999999 |
|  |  | *100* | 1.35 | 0.73 |  | 0.14 | 0.1 |  | 0.16 |
|  |  | *150* | 3.92 | 0.96 |  | 1.43 | 0.6 |  | 0.06 |
|  |  | *200* | 6.35 | 1.50 |  | 3.79 | 1.06 |  | 0.24 |
|  |  | *250* | 9 | 1.60 |  | 6.86 | 1.23 |  | 0.32 |
|  |  | *300* | 12.07 | 1.56 |  | 9.79 | 1.31 |  | 0.27 |
|  |  | *350* | 14.64 | 1.40 |  | 12.29 | 1.41 |  | 0.28 |
|  |  | *400* | 17 | 1.27 |  | 14 | 1.51 |  | 0.27 |
|  |  | *450* | 18.42 | 1.20 |  | 15.86 | 1.61 |  | 0.38 |
|  |  | *500* | 19.5 | 1.12 |  | 17.5 | 1.44 |  | 0.31 |
|  |  | *550* | 21 | 1.14 |  | 19.21 | 1.34 |  | 0.46 |
|  |  | *600* | 21 | 1.26 |  | 20.21 | 1.2 |  | 0.78 |
|  | **pIC** | *0* | 0 | 0 | 20/15 | 0 | 0 | 16/12 | >0.999999 |
|  |  | *50* | 1.05 | 0.42 |  | 0.94 | 0.48 |  | >0.999999 |
|  |  | *100* | 4.75 | 1.18 |  | 3.13 | 1.03 |  | 0.49 |
|  |  | *150* | 9.95 | 1.57 |  | 6.5 | 1.44 |  | 0.13 |
|  |  | *200* | 14.95 | 1.78 |  | 9.81 | 1.64 |  | 0.09 |
|  |  | *250* | 19.15 | 1.95 |  | 13.25 | 1.47 |  | 0.06 |
|  |  | *300* | 21.95 | 1.97 |  | 15.88 | 1.33 |  | 0.06 |
|  |  | *350* | 23.25 | 2.14 |  | 18.06 | 1.27 |  | 0.15 |
|  |  | *400* | 23.45 | 2.14 |  | 19.63 | 1.2 |  | 0.26 |
|  |  | *450* | 23 | 2.42 |  | 20.94 | 1.09 |  | 0.39 |
|  |  | *500* | 22.85 | 2.45 |  | 21.38 | 0.9 |  | 0.52 |
|  |  | *550* | 22.85 | 2.71 |  | 21.63 | 1.14 |  | 0.75 |
|  |  | *600* | 19.35 | 3.14 |  | 16.88 | 1.88 |  | 0.7 |
