## Supplementary material for "Sexual differences in neuronal and synaptic properties across subregions of the mouse insular cortex": sup table 2

**Supplementary Table 2. Quantitative evaluation of amplitude, frequency, and kinetics of excitatory and inhibitory events in adult male and female of anterior and posterior IC neurons**

| **Misure** | **Area** | **Sex** | **Median** | **MAX** | **MIN** | **n/N** | ***Two-way Anova Main effects and interactions*** | ***Multiple comparison (Šidák’s post hoc test)*** |
| --- | --- | --- | --- | --- | --- | --- | --- | --- |
| **sEPSCs Amplitude (pA)** | aIC | M | 13.71 | 18.09 | 11.82 | 14/9 | Interaction: p = 0.6224 Area:  F (1. 67) = 1.763 p = 0.1888 Sex:  F (1. 67) = 1.892 p = 0.1735 | aIC Male vs pIC Male  p = 0.0019  aIC Female vs pIC Female p = 0.0241  aIC Male vs aIC Female p = 0.3748  pIC Male vs pIC Female p = 0.0480 |
|  |  | F | 14.95 | 19.09 | 10.24 | 13/6 |  |  |
|  | pIC | M | 16.33 | 24.25 | 11.12 | 27/16 |  |  |
|  |  | F | 16.08 | 20.81 | 11.33 | 17/12 |  |  |
| **sEPSCs AFrequency (Hz)** | aIC | M | 4.445 | 6.53 | 1.77 | 14/9 | Interaction: p = 0.0038 Area:  F (1. 66) = 0.3349 p = 0.5648 Sex:  F (1. 66) = 1.613 p = 0.2086 | aIC Male vs pIC Male  p = 0.0187  aIC Female vs pIC Female p = 0.2042  aIC Male vs aIC Female p = 0.4204  pIC Male vs pIC Female p = 0.0054 |
|  |  | F | 3.68 | 6.22 | 1.04 | 13/6 |  |  |
|  | pIC | M | 2.235 | 6.17 | 0.83 | 27/16 |  |  |
|  |  | F | 4.63 | 7.99 | 0.49 | 17/12 |  |  |
| **sEPSCs Rise (ms)** | aIC | M | 0.675 | 0.86 | 0.55 | 14/9 | Interaction: p = 0.6224 Area:  F (1. 67) = 1.763 p = 0.1888 Sex:  F (1. 67) = 1.892 p = 0.1735 | aIC Male vs pIC Male  p = 0.7830  aIC Female vs pIC Female p = 0.4008  aIC Male vs aIC Female p = 0.7961  pIC Male vs pIC Female p = 0.3203 |
|  |  | F | 0.7 | 0.85 | 0.35 | 13/6 |  |  |
|  | pIC | M | 0.66 | 0.97 | 0.41 | 27/16 |  |  |
|  |  | F | 0.6 | 0.84 | 0.4 | 17/12 |  |  |
| **sEPSCs Decay (ms)** | aIC | M | 6.63 | 9.41 | 5.05 | 14/9 | Interaction: p = 0.6530 Area:  F (1. 61) = 0.1419 p = 0.7077 Sex:  F (1. 61) = 1.072 p = 0.3046 | aIC Male vs pIC Male  p = 0.7966  aIC Female vs pIC Female p = 0.9983  aIC Male vs aIC Female p = 0.5384  pIC Male vs pIC Female p = 0.8875 |
|  |  | F | 6.055 | 10.37 | 0.45 | 13/6 |  |  |
|  | pIC | M | 6.56 | 10.1 | 0.47 | 27/16 |  |  |
|  |  | F | 6.01 | 11.94 | 2.06 | 17/12 |  |  |
| **sIPSCs Amplitude (pA)** | aIC | M | 38.83 | 68.11 | 18.89 | 22/10 | Interaction: p = 0.4648 Area:  F (1. 68) = 2.933 p = 0.0913 Sex:  F (1. 68) = 2.380 p = 0.1275 | aIC Male vs pIC Male  p = 0.7441  aIC Female vs pIC Female p = 0.1651  aIC Male vs aIC Female p = 0.2018  pIC Male vs pIC Female p = 0.8198 |
|  |  | F | 44.53 | 64.1 | 23.39 | 15/10 |  |  |
|  | pIC | M | 35.91 | 62.49 | 18.36 | 14/11 |  |  |
|  |  | F | 34.58 | 67 | 18.89 | 19/11 |  |  |
| **sIPSCs Frequency (Hz)** | aIC | M | 5.17 | 8.32 | 1.75 | 22/10 | Interaction: p = 0.0604 Area:  F (1. 68) = 1.589 p = 0.2118 Sex:  F (1. 68) = 0.8139 p = 0.3702 | aIC Male vs pIC Male  p = 0.0587  aIC Female vs pIC Female p = 0.8735  aIC Male vs aIC Female p = 0.7327  pIC Male vs pIC Female p = 0.0948 |
|  |  | F | 4.12 | 8.05 | 1.88 | 15/10 |  |  |
|  | pIC | M | 2.93 | 6.54 | 1 | 14/11 |  |  |
|  |  | F | 4.39 | 10.08 | 0.62 | 19/11 |  |  |
| **sIPSCs Rise (ms)** | aIC | M | 0.55 | 0.85 | 0.33 | 14/10 | Interaction: p = 0.3286 Area:  F (1. 61) = 3.055 p = 0.0855 Sex:  F (1. 61) = 0.2303 p = 0.6330 | aIC Male vs pIC Male  p = 0.1341  aIC Female vs pIC Female p =0.8177  aIC Male vs aIC Female p = 0.9173  pIC Male vs pIC Female p = 0.5387 |
|  |  | F | 0,565 | 0.68 | 0.5 | 18/10 |  |  |
|  | pIC | M | 0.605 | 0.9 | 0.52 | 27/10 |  |  |
|  |  | F | 0.58 | 0.97 | 0.39 | 13/10 |  |  |
| **sIPSCs Decay (ms)** | aIC | M | 7.295 | 9.37 | 5.24 | 16/10 | Interaction: p = 0.9436 Area:  F (1. 55) = 8.056 p = 0.0063 Sex:  F (1. 55) = 0.8569 p = 0.3586 | aIC Male vs pIC Male  p = 0.0960  aIC Female vs pIC Female p =0.0974  aIC Male vs aIC Female p = 0.6986  pIC Male vs pIC Female p = 0.8180 |
|  |  | F | 7.41 | 8.01 | 2.96 | 14/01 |  |  |
|  | pIC | M | 8.955 | 11.09 | 4.92 | 19/10 |  |  |
|  |  | F | 7.63 | 10.7 | 5.22 | 13/10 |  |  |
